# α-Helix stabilization by co-operative side chain charge-reinforced interactions to phosphoserine in a basic kinase-substrate motif

**DOI:** 10.1101/2021.11.27.470016

**Authors:** Matthew Batchelor, Robert S. Dawber, Andrew J. Wilson, Richard Bayliss

## Abstract

How cellular functions are regulated through protein phosphorylation events that promote or inhibit protein–protein interactions (PPIs) is key to understanding regulatory molecular mechanisms. Whilst phosphorylation can orthosterically or allosterically influence protein recognition, phospho-driven changes in the conformation of recognition motifs are less well explored. We recently discovered that clathrin heavy chain recognises phosphorylated TACC3 through a helical motif that, in the unphosphorylated protein, is disordered. However, it was unclear whether and how phosphorylation could stabilize a helix in a broader context. In the current manuscript, we address this challenge using poly-Ala based model peptides and a suite of circular dichroism and nuclear magnetic resonance spectroscopies. We show that phosphorylation of a Ser residue stabilizes the α-helix in the context of an Arg_(*i* – 3)_pSer_*i*_ Lys_(*i* + 4)_ triad through charge-reinforced side chain interactions with positive co-operativity, whilst phosphorylation of Thr induces an opposing response. This is significant as it may represent a general method for control of PPIs by phosphorylation; basic kinase-substrate motifs are common with 55 human protein kinases recognising an Arg at a position –3 from the phosphorylated Ser, whilst the Arg_(*i* – 3)_pSer_*i*_ Lys_(*i* + 4)_ is a motif found in over 2000 human proteins.

## Introduction

The determinants of α-helix stability are key in understanding the formation and strength of α-helix mediated protein-protein interactions (PPIs),^1^ and therefore in enabling peptide drug-discovery.^2, 3^ Prior studies established helix propensities of individual amino acids,^4, 5^ the role of helix capping^6, 7^ and effects of interaction between side chains,^8–11^ however the role of phosphorylation is less well explored.

Classically, phosphorylation occurs within solvent-exposed loops or disordered regions.^12^ Such phosphorylation events play a regulatory role in the PPI network of substrate proteins; primed with a new highly charged phosphate group as a recognition feature, such regions can bind to a different protein to activate or deactivate a subsequent cellular process.^13^ More recently, phosphorylation has been shown to affect secondary/tertiary structure, *e.g*., the intrinsically disordered 4E-BP2 undergoes a disorder-to-helix transition upon binding to eIF4E, but on multisite phosphorylation, folds instead into a four-stranded β-domain.^14^ Ser phosphorylation has been shown to increase helicity in model peptides,^15^ notably at the N-terminus,^16, 17^ a feature exploited in design of phosphorylation-stabilized tertiary helix structures.^18, 19^ Secondary structure has also been shown to be destabilized by phosphorylation^20, 21^ with internal phosphorylation of Thr and Ser residues helix destabilizing.^15–17^ Herein we describe the effects of phosphorylation on helix stability using model poly-Ala based peptides, into which we grafted a motif inspired by studies on the Aurora A/TACC3/CHC pathway, an unusual example of a protein–protein interaction centred on a phosphorylated helix.^22^

Aurora A-mediated phosphorylation of TACC3 promotes complex formation with clathrin heavy chain (CHC) protein resulting in molecular ‘bridges’ across parallel microtubules to ensure accurate chromosome alignment and segregation during mitosis. The TACC3/CHC complex is α-helix-mediated;^22^ the TACC3 residue phosphorylated by Aurora A, Ser558, is in a basic kinase-substrate motif, flanked by basic Arg555 and Lys562 residues in *i* – 3 and *i* + 4 positions rendering them suitably placed to stabilize helicity *via* interactions with the phosphate (Fig. 1). The TACC3-inspired R_(*i* − 3)_pS_*i*_K_(*i* + 4)_ motif was grafted into a well-studied poly-Ala-based model peptide.^23, 24^ Poly-Ala peptides are good models due to their intrinsic α-helix-forming propensity; sequence modifications can be related to the introduction/deletion of potential interactions between side chains.^9, 25^ Peptides were designed to investigate the α-helix stabilizing influence of phosphate salt bridging. The phosphorylation state was altered, Arg and Lys side chains were systematically moved in and out of pSer registry, and the effect of substitution of Ser for Thr was studied.

**Figure 1.**
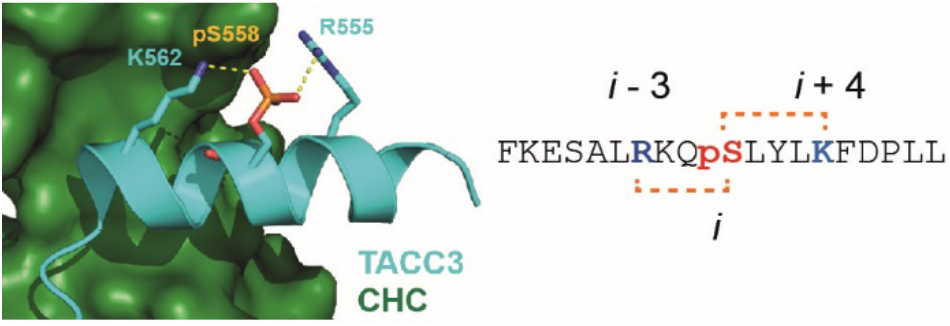
X-Ray crystal structure for the TACC3/CHC interaction (PDB ID: 5ODS; TACC3 cyan, CHC forest green), together with the TACC3 sequence; the R_(*i* – 3)_pS_*i*_K_(*i* + 4)_ motif is highlighted with side chains shown as sticks.

## Experimental Procedures

### Synthesis and Purification of Peptides

Amino acids were purchased from Novabiochem (Merck), Sigma-Aldrich or Fluorochem. All amino acids were N-Fmoc protected and side chains were protected with Boc (Lys), ^t^Bu (Ser, Thr), Trt (Gln) or Pbf (Arg). Solvents and reagents used in peptide synthesis were sourced as follows: DMF and DCM (ACS grade) from Sigma-Aldrich; piperidine (99%) from Alfa Aesar; DIPEA (reagent grade) from Fluorochem; TFA (peptide grade) from Fluorochem; acetic anhydride (>99%) from Sigma Aldrich; HCTU (reagent grade) from Fluorochem; formic acid (analytical reagent grade) from Fisher Scientific; *m*-Cresol (99%) from Acros Organics; thioanisole (99%) from Alfa Aesar; EDT (95%) from Sigma Aldrich. Peptides were synthesized on Rink Amide MBHA resin by room temperature Fmoc-Solid Phase Peptide Synthesis (SPPS) using the following cycle on an automated peptide synthesizer (CEM Liberty Blue). *Deprotection* – clean resin dip tube, wash with DMF (15 mL), add DMF: piperidine: formic acid (75:20:5) solution (6 mL), room temperature method (5 min), wash with DMF (15 mL), add DMF: piperidine: formic acid (75:20:5) solution (6 mL), room temperature method (5 min), wash with DMF (15 mL), clean resin dip tube, wash with DMF (15 mL). *Coupling* – add amino acid solution (0.2 M, 2.5 mL), add coupling reagent (HCTU; 0.2 M, 1 mL), add activator base (DIPEA; 0.2 M, 0.5 mL), room temperature method (18 min), wash with DMF (15 mL), drain. For double couplings, this step was repeated. *N-terminal acetylation* – after coupling of the final residue, the resin was ejected from the reaction vessel and N-terminal acetylation/cleavage and deprotection was performed manually. Acetic anhydride (10 *eq.*) and DIPEA (10 *eq.*) were dissolved in DMF (3 mL) and the solution was transferred to the resin. After 2 h, the resin was drained, washed with DMF (3 × 2 mL × 2 min) and successful capping determined by a negative color test (Kaiser test).^26^ *Cleavage and deprotection* – the resin was washed with DMF, DCM and Et2O (for each: 3 × 3 mL × 2 min). Peptides were then simultaneously cleaved and side chain deprotected with a prepared Reagent K cleavage cocktail (6 mL): TFA/m-cresol/H_2_O/thioanisole/EDT (82.5/5/5/5/2.5). After 3 h, the resin was washed with fresh TFA (3 mL × 2 min) and the solution concentrated under a flow of N_2_. The resulting oil was precipitated with ice-cold Et_2_O (10 mL) and placed in a centrifuge (3000 rpm × 3 min). The supernatants were removed, the precipitate rinsed with ice-cold Et_2_O (3 × 10 mL) and dried under a flow of N_2_. *Purification* – peptides were purified by preparative HPLC using a Kinetex 5 μM EVO C18 preparative column (reversed phase) on an increasing gradient of acetonitrile to water (plus 0.1% TFA v/v in water) at a flow rate of 10 mL min^−1^. Crude peptides were dissolved in minimal amounts of acetonitrile: water (1:1). Purification runs injected a maximum of 5 mL of crude peptide solution and were allowed to run for 35 min, with acetonitrile increasing from 5 to 95%, and the eluent scanned with a diode array at 210, 254 and 280 nm. Fractions were checked by LCMS, collected and lyophilized. Final purity of peptides was confirmed by high-resolution mass spectrometry and analytical HPLC (Supplementary Information, Table S1).

### Peptide solutions

‘CD buffer’ containing 1 mM sodium borate, 1 mM sodium citrate, 1 mM sodium phosphate, and 10 mM NaCl, pH 7,^15^ was used to facilitate easy alteration of peptide solution pH through addition of small amounts of HCl or NaOH. This buffer has low absorbance in the region of wavelengths (190–260 nm) used for CD measurements. Solid lyophilized peptides were dissolved to give 1–2 mM solutions based on mass using CD buffer resulting in mildly acidic solutions (pH ~ 6–7). Solution absorbance measurements at 280 nm were used to calculate more accurately the peptide concentration, based on a Tyr extinction coefficient of 1280 M^−1^ cm^−1^,^27, 28^ with 1 mM concentrations based on mass typically resulting in ~0.7–1.0 mM solutions based on absorbance. Initial (~1 mL) sample pH measurements were made using a three-point-calibrated meter equipped with a microelectrode probe. *In situ* pH measurements (*e.g*. during CD experiments) were made by careful removal of small volumes of sample to test with pH strips. *In situ* estimates of pH during NMR experiments were made using the buffer phosphate chemical shift by comparison with CD buffer-only samples calibrated at pH 5, 6, 7 and 8.

### CD measurements

CD spectra were recorded using an APP Chirascan CD spectropolarimeter and 1 mm pathlength quartz cuvettes. Sample concentrations were typically 50–100 μM and once diluted to this degree had a pH matching the buffer alone (pH 7). Spectra were recorded at 5 °C over wavelengths ranging from 260 to 180 nm, with data collected every 1 nm (1 nm/s), and each spectrum was recorded in duplicate. Temperature ramping experiments were recorded from 5 to 70–80 °C with data recorded every ~1 °C using a settling time between measurements of 120 s. A background spectrum for CD buffer alone was recorded for each data collection session, and background ellipticity values were subtracted from raw sample ellipticity values (*θ*) when calculating mean residue ellipticities (MREs) using:

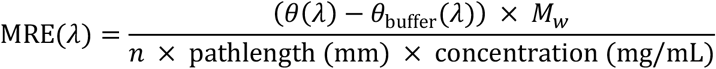

where *M_w_* is the peptide molecule weight, and *n* is the number of total peptide bonds, in each case taken to be 16 accounting for the acetyl capping group.

Several methods are available to estimate the helicity based on MRE values.^29–31^ Estimates of peptide helicity were made using the relation used by Baker *et al*.^32^ as follows:

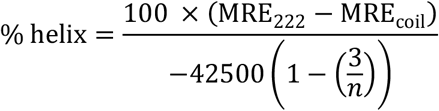

where MRE_222_ is the MRE value at 222 nm, MRE_coil_ = 640 − 45*T* (with *T* in °C) = 415 deg cm^2^ dmol^−1^ res^−1^ at 5 °C and *n* is the number of backbone amide bonds including the *N*-terminal acetyl (as above). For a selection of peptides, MRE values were calculated from data collected over a wider (100x) range of concentrations (roughly 7–700 μM, Fig. S1). No change in MRE values was observed suggesting that peptide self-association does not occur.

### Analysis of energetics

A simple estimate of the Gibbs free energy change (Δ*G*) of the helical (α) peptide in solution relative to its unfolded (u) state, can be calculated under a two-state, all-or-nothing approximation from the % helicity values determined experimentally by CD spectroscopy.^11^ The equilibrium constant *K*_α_ between the folded and unfolded conformations in solution is defined by:

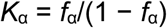

where *f*α is the fraction of helical and (1 − *f*_α_) = *f*_u_, the fraction of unfolded peptide. The Gibbs free energy change on going from the unfolded state to the helical state is:

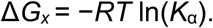

The change in the Gibbs free energy change on folding (ΔΔ*G*) is calculated relative to the least helical pSer containing peptide **3** (ΔΔ*G* = Δ*G_x_* – Δ*G*_**3**_) to highlight how the introduction of potential sites for non-covalent interaction into the sequence influences conformational energetics; the *i* + 5 spacings in peptide **3** mean that it has no potential for inter-side-chain interaction for a helical conformation.

### NMR experiments

NMR spectra were recorded on a 600-MHz Bruker Avance spectrometer equipped with a quadruple resonance QCI-P cryoprobe, or a 950-MHz Bruker Ascend Aeon spectrometer equipped with a 5 mm TXO cryoprobe. For each peptide, natural abundance ^1^H–^15^N HMQC and ^1^H–^13^C HSQC (constant-time) spectra were recorded for 0.5–1.0 mM samples at 5 °C. The low temperature maximizes the helical content of each peptide, and also (when combined with using pH between 6 and 7) slows proton exchange enough to allow the Arg Hε–Nε correlation to be visible. Clean spectra ^1^H–^15^N HMQC took ~12 h to record. To reduce experiment time (to ~ 6 h), ^1^H–^13^C HSQC spectra were recorded with reduced spectral width in ^13^C (typically 20 ppm centred on 56 ppm), and the spectrum folded out to show the position of the non-Hα–Cα correlations. ^1^H–^1^H TOCSY and ^1^H–^1^H NOESY spectra were recorded and used to assign resonances for each peptide through residue-specific shifts and strong (*i*, *i* – 1) NOE connectivity. Spectra were initially processed using NMRPipe/NMRDraw,^33^ and resonance assignments were carried out using CCPNmr Analysis.^34^ Systematic patterns of peak shifts in spectra for different but related peptides were used to aid assignment. Full assignment of backbone resonances was possible (with the exception of uncharacterized amide carbonyl carbons). ^13^C resonance assignment in ^1^H–^13^C HSQC spectra required a unique attached ^1^H resonance and/or the peak position to occur in a residue-specific region of the spectrum. In a few instances, Ala Cα and Cβ resonances could not be independently assigned so two positions—marking the potential range in Cα/Cβ shift—were recorded. Secondary shifts (Δδ) = observed shifts (δ) – expected shifts for that residue in a random coil conformation (δ_RC_), were calculated using recently published δ_RC_ values for phosphorylated residues^35^ available via the website: (https://www1.bio.ku.dk/english/research/bms/sbinlab/randomchemicalshifts2/). 1D ^31^P spectra were recorded on the 600-MHz spectrometer using the 30-degree flip angle pulse sequence.

## Results

### Basic residues in flanking positions enhance the helicity of peptides incorporating a pSer residue

A range of peptides (**1**–**10**, Table 1) were designed and prepared using solid-phase synthesis as *N-*terminal acetamides and *C-*terminal amides (see Supplementary Information for analytical characterization data). The relative helical propensity for each of the peptides was established using far UV circular dichroism (CD) spectroscopy.^27^ Fig. 2a compares the CD spectrum of the phosphorylated and unphosphorylated peptides **1** and **2**. **1** exhibits a CD profile characteristic of an α-helix with negative bands at 222 nm and 208 nm, and a positive band at ~190 nm (Fig. 2a, see Supplementary Information Fig. S1 for data demonstrating spectra are concentration independent). For peptide **2**, reduced helicity is indicated from smaller bands at 222 and 190 nm, and a shift of the peak at 208 nm to lower wavelength. Estimation of the % helicity using the MRE value at 222 nm gave 29% for **1** and 22% for **2** (Table 1), indicating that phosphorylation enhances helicity with *i* – 3 Arg and *i* + 4 Lys basic side chains. Variant pSer to Ala control peptides were more helical as would be expected given the increased helical propensity of Ala (peptides **9** and **10**, Table 1, Fig. S2).^4^ The variation in MRE222 with temperature indicated a broad transition from part-helical to coil structures (all peptides converged at ~4000 deg cm^2^ dmol^−1^ at ~70 °C) and weak folding co-operativity as expected for short peptides of this length (Fig. S3).

**Table 1.**
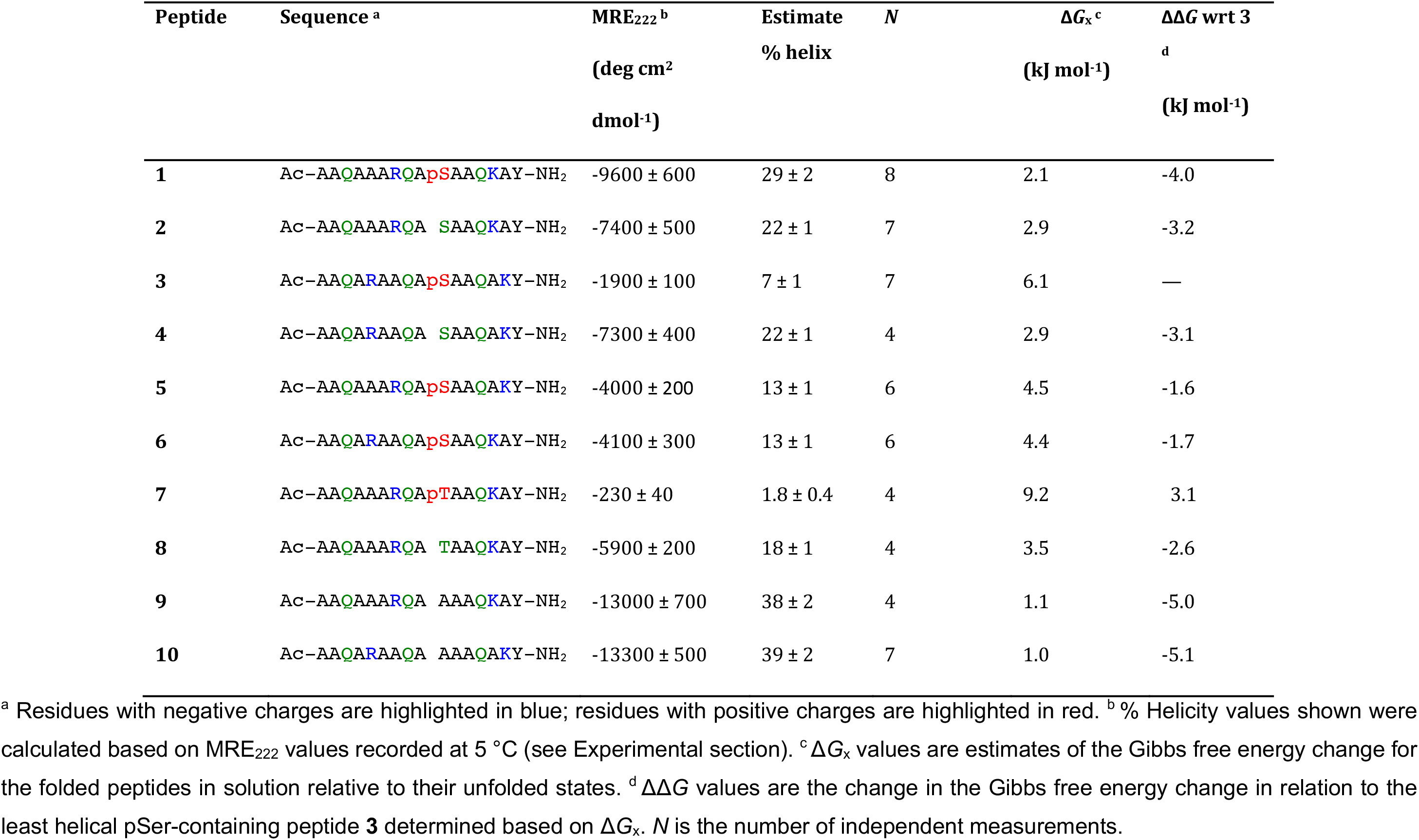
The sequences for poly-Ala model peptides **1–10** prepared and analysed in this study.

**Figure 2.**
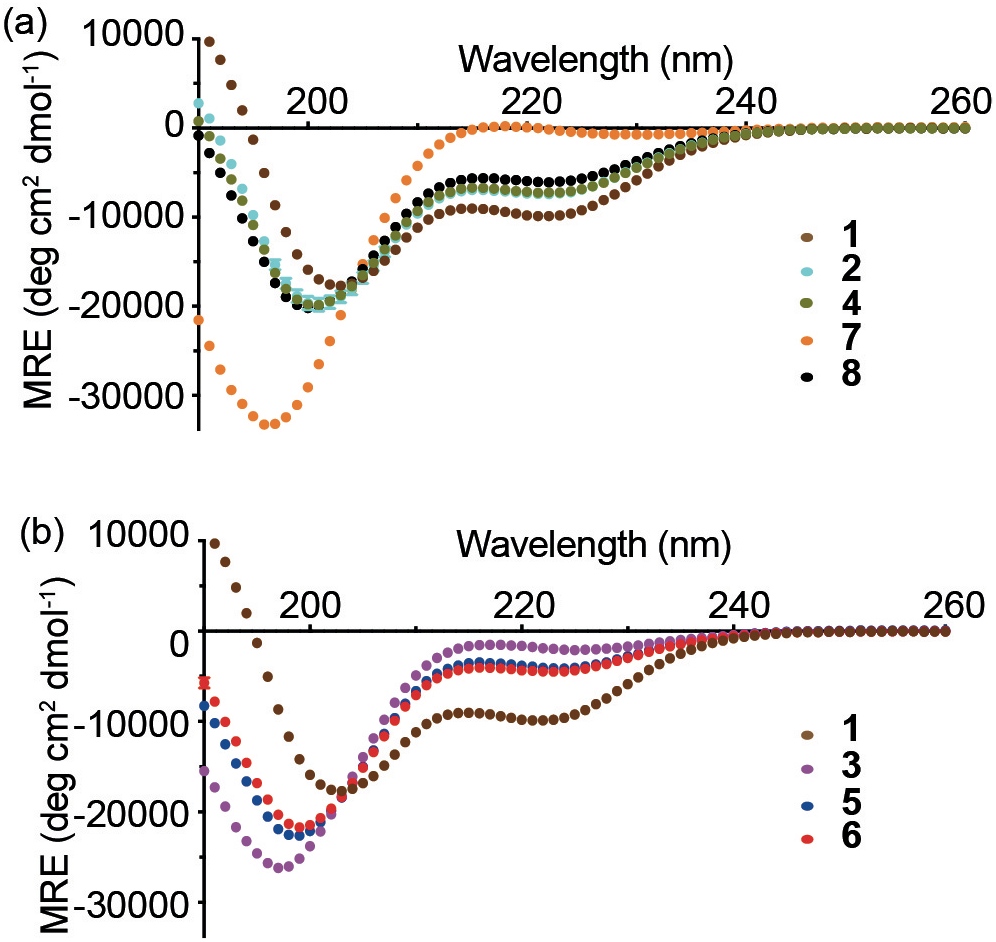
CD spectroscopy for model peptides. (a) CD spectra for peptides **1–2** and **7–8** comparing the contrasting effect of phosphorylation on the R_(*i* – 3)_S_*i*_K_(*i* + 4)_ and R_(*i* – 3)_T_*i*_K_(*i* + 4)_ motifs. Also shown is the spectrum for **4**, with its R_(*i* – 5)_S_*i*_K_(*i* + 5)_ motif. (b) CD spectra for **1**, **3, 5** and **6** illustrating the effect of moving one or two basic groups out of ‘helical-registry’ with the pSer. Experiments were carried out at 5 °C in CD buffer (10 mM NaCl, 1 mM sodium phosphate, 1 mM sodium borate, and 1 mM sodium citrate) at pH 7. Peptide concentrations were in the range 50–100 μM.

The effect on peptide helicity of moving Arg and Lys out of helical-registry was explored (Fig. 2b). In a peptide where no helix-compatible side-chain interactions with the phosphate are possible, α-helicity was lowest (**3**, 7%), as expected given the propensity of pSer to be destabilizing in the centre of an α-helix.^17^ Indeed, the unphosphorylated version (**4**) has higher helicity, matching that of peptide **2**. The similarity of the helicity of peptides **4** and **2** (and peptides **9** and **10**) indicates that the position of Lys and Arg has little effect on the overall helicity when there is no phosphoryl group with which to interact. Having an Arg residue *i* – 3 or Lys residue *i* + 4 to the pSer increased the α-helix-forming propensity (**5** and **6**, both 13%). CD analyses on the pThr-containing peptide **7** indicated low helicity (2%) whereas the Thr analogue elicited higher helicity (**8**, 18%) similar to unphosphorylated Ser peptide **2** (Fig. 2a). This can be attributed to the proclivity of pThr to adopt a compact conformation that is incompatible with a helical conformation and is evidently not countered by the potential benefit of non-covalent interaction with side chains as for pSer (peptide **1**).^36^

### Side chain interactions that enhance pSer peptide helicity are pH dependent and positively cooperative

The effect of pH on helicity for peptide **1** provided useful insights on the contribution of charge-based interactions between side chains (Fig. 3). Protonating the phosphate by reducing the pH below the p*K*_a_ of PO_3_^2−^ (pH < 5 forming singly-charged PO_3_H^−^) or deprotonating the Lys or Arg groups at high pH would be anticipated to reduce or eliminate charge-reinforced interactions between side chains. The estimated helicity of peptide **1** at low pH is ~16% and at high pH is ~6%, similar values to those of peptide **5**/**6** (with a single potential X^+^–PO_3_^2−^ interaction) and peptide **3** (no interaction), respectively. For comparison, the more limited effect of changing pH on peptide helicity for other selected peptides is shown in the Supplementary Information (Fig. S4). We used the measured % helicity values to estimate the Gibbs free energy change of the α-helical peptides in solution relative to their unfolded states, Δ*G* (Table 1).^11^ This allowed us to make quantitative estimates of the change in the Gibbs free energy change, ΔΔ*G*, associated with formation of charge-reinforced side chain interactions involving the phosphate. By relating the Δ*G* of **5** to that of **3**, the formation of such an interaction between the phosphate group and *i* – 3 Arg side chain has a ΔΔ*G* of −1.6 kJ mol^−1^. Comparison of Δ*G* values for **6** and **3** show that potential interaction between the phosphate and the *i* + 4 Lys side chain has a similar ΔΔ*G* (−1.7 kJ mol^−1^). These values are in agreement with typical values for formation of charge-reinforced interactions between side chains in helices,^37, 38^ *e.g*. Glu–Lys (*i*, *i* ± 3 or *i*, *i* ± 4) interactions under similar conditions gave rise to ΔΔ*G* values of 1.2–1.7 kJ mol^−1^.^38^ For both **5** and **6**, the ability to form interactions between side chains is insufficient to offset the destabilizing effect of introducing a pSer in the middle of the helical sequence (**4** *vs*. **3**, ΔΔ*G* = −3.1 kJ mol^−1^).^16^ In contrast, **1**, where both interactions can form, would be expected to have a reported ΔΔ*G* (relative to **3**) of approximately −3.3 kJ mol^−1^ if the contributions were additive (only marginally higher than the ΔΔ*G* for **2**). However, with a ΔΔ*G* of −4.0 kJ mol^−1^, this indicates that the effects of two charge-reinforced interactions with phosphate are positively co-operative.

**Figure 3.**
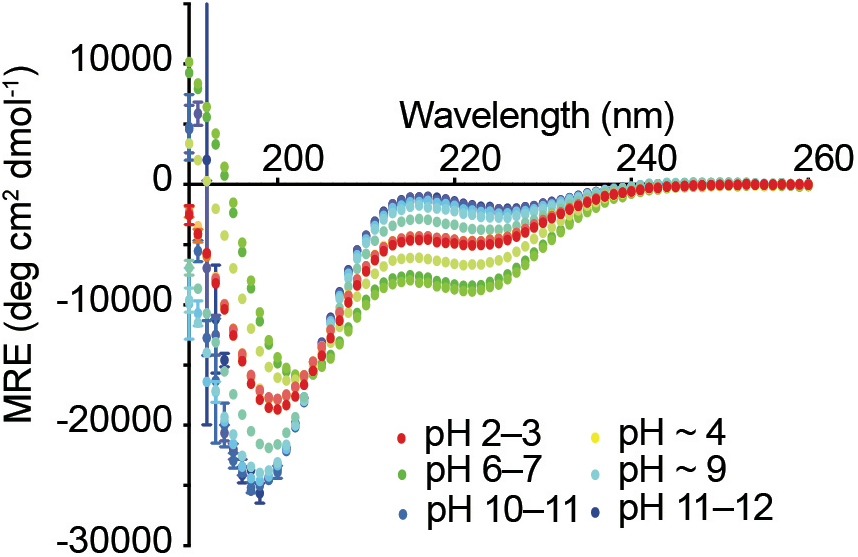
The effect of pH on the helicity of peptide **1**. Low and high pH reduce helicity indicating that charge-reinforced side chain interactions stabilize the helix. Experiments were carried out at 5 °C in CD buffer (10 mM NaCl, 1 mM sodium phosphate, 1 mM sodium borate, and 1 mM sodium citrate). Peptide concentrations were in the range 50–100 μM. The pH was altered by addition of small amounts of 0.1 M HCl or 0.05 M NaOH, with volume changes factored-in to concentration and MRE calculations.

### NMR chemical shifts show increased helicity in pSer peptides with in-helical-range basic residues

To provide additional structural insight to complement the CD analyses, a suite of four NMR spectra was recorded for each peptide to assign ^1^H, ^13^C and ^15^N resonances. Residue assignments in a ^1^H–^15^N HMQC spectrum were achieved using a combination of ^1^H–^1^H TOCSY/NOESY spectra, informed by a ^1^H–^13^C HSQC spectrum and systematic peak position shifts between spectra for different peptides. Natural abundance ^1^H–^15^N HMQC spectra for peptides **1** and **2** are shown in Fig. 4a; phosphorylation leaves H–N peak positions for residues 1–5 largely unaffected whilst a characteristic downfield shift is observed for the backbone H–N of pSer with respect to Ser as marked by the red line.^39^ Neighbouring residues also exhibit shifts with the phosphorylated peptide **1** showing greater peak dispersion than the unphosphorylated peptide **2**, suggesting increased secondary structure content. Patterns in Cα shifts also provide a useful guide to local secondary structure. Fig. 4b shows the Hα–Cα region of ^1^H–^13^C HSQC spectra for two examples, peptide **1** and peptide **3** with Arg and Lys in- and out-of-helical-range, respectively. Systematic downfield Cα shifts and upfield Hα shifts observed for peptide **1** compared to peptide **3** indicate increased helical content. To account for changes in sequence between peptides, secondary shifts (Δδ) (observed shifts (δ) – expected shifts for that residue in a random coil conformation (δ_RC_)) ^40–42^ were calculated using random coil values that were recently updated to include values for phosphorylated residues (Fig. 5).^35^ Data for peptides **4**, **9** and **10** are shown in the Supplementary Information, Fig. S5. These data largely mirror the CD data and show the weighting of the more helical residues towards the N-terminus. It was noted that the pThr Cα secondary shift values are higher than expected for a residue in a random coil configuration, indicating that pThr occupies an arrangement unlike that in the QQpTQQ peptide used to generate the values.^35^ Example spectra showing the expected effect of pH on pSer and pThr Hα–Cα correlations are given in the Supplementary Information (Fig. S6).

**Figure 4.**
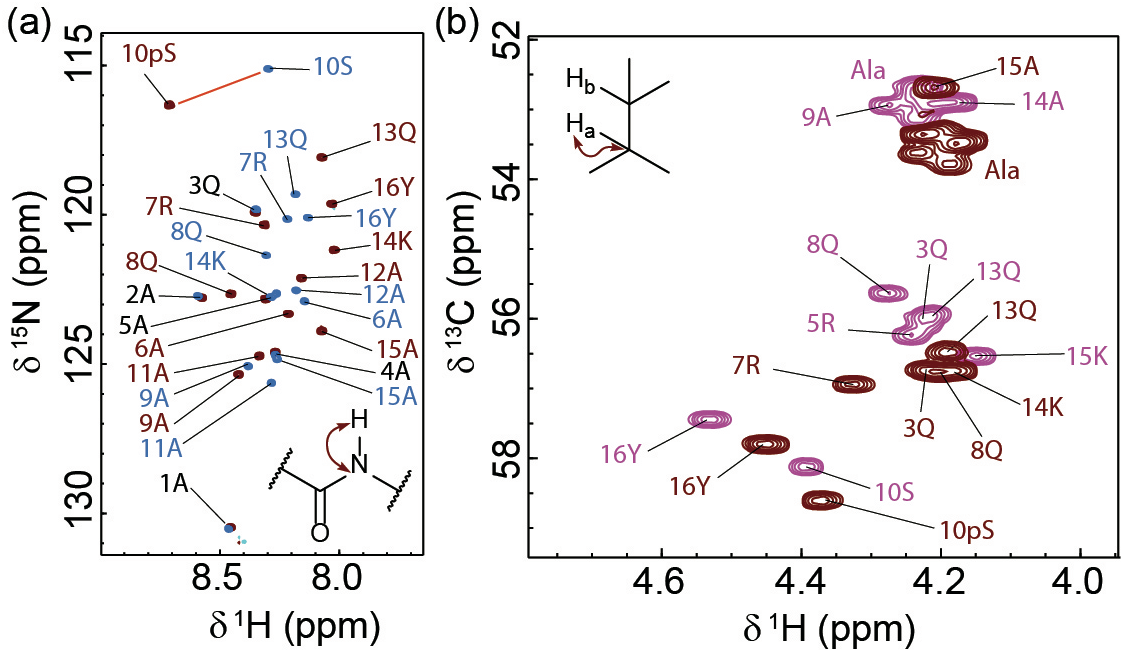
NMR spectroscopy of peptides, (a) comparison of part of the ^1^H–^15^N HMQC spectra for peptides **1** (brown) and **2** (blue), showing the effect of phosphorylation on main-chain HN peaks of the R_(*i* – 3)_S_*i*_K_(*i* + 4)_ motif peptide; (b) ^1^H–^13^C HSQC spectra for peptides **1** (brown) and **3** (magenta). These spectra highlight the difference between a phospho-peptide that is capable (**1**) or incapable (**3**) of forming inter-turn interactions between the phosphate group and neighbouring basic residues (Arg and Lys). Spectra were recorded on a 600-MHz Bruker Avance spectrometer equipped with a quadruple resonance QCI-P cryoprobe, or a 950-MHz Bruker Ascend Aeon spectrometer equipped with a 5 mm TXO cryoprobe. Experiments were performed in CD buffer (10 mM NaCl, 1 mM sodium phosphate, 1 mM sodium borate, and 1 mM sodium citrate, pH 6.5–7) at 5 °C. Peptide concentrations were in the range 0.5–1.0 mM

**Figure 5.**
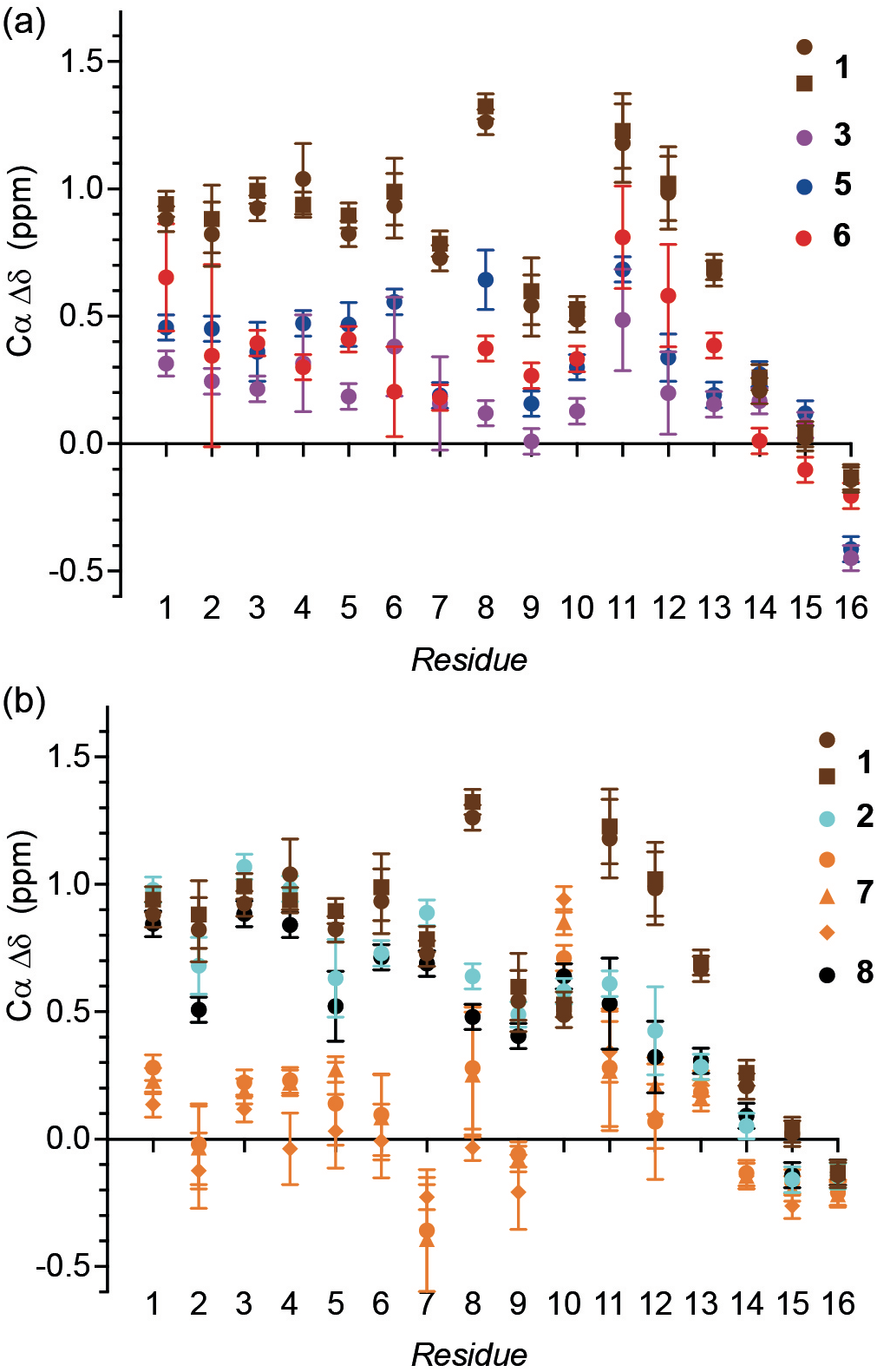
(a, b) Residue specific Cα secondary shifts at 5 °C and pH 6.5–7 are plotted for each residue for peptides studied **1**–**8**. Different values shown for peptide **1** and **7** represent measurements carried out at pH values between 6 and 7.5, without significant variation. Spectra were recorded on a 600-MHz Bruker Avance spectrometer equipped with a quadruple resonance QCI-P cryoprobe, or a 950-MHz Bruker Ascend Aeon spectrometer equipped with a 5 mm TXO cryoprobe. Experiments were performed in CD buffer (10 mM NaCl, 1 mM sodium phosphate, 1 mM sodium borate, and 1 mM sodium citrate). The pH was altered by addition of small amounts of 0.1 M HCl or 0.05 M NaOH. Peptide concentrations were in the range 0.5–1.0 mM.

### Interaction effects shown through specific side chain resonance analyses

Confirmation of intra-helix side chain interactions was obtained using additional NMR experiments focussed on side chain resonances. Performing experiments at a low temperature of 5 °C maximizes the helical content and when coupled with use of low pH (below pH 7) also reduces proton exchange enough for Arg Hε–Nε side chain groups to be visible in ^1^H–^15^N HMQC spectra.^43^ Each peptide has a single Arg residue resulting in a single Hε–Nε correlation (Fig. 6a). The Arg Hε shift appears at ~7.25 ppm for all peptides with the exception of peptide **5** (7.5 ppm) and peptide **1** (7.8 ppm), both of which have Arg in the *i* – 3 position relative to the phosphorylated residue—a position suitable for interactions between side chains across a helical turn—*and* have been shown to be helical. The magnitude of the downfield shift with respect to peptides that have no potential to form such interactions correlates with the overall helicity of the peptide and thus peptide **1** has the largest downfield shift, peptide **5** has an intermediate value and peptide **7** is unmoved. The Hε shift for peptide **1** is similar to those reported for Arg residues in a single-alpha helix domain that had three potential Glu salt bridge partners (7.7–7.9 ppm).^44^ In line with the CD analyses, on decreasing pH, the Hε–Nε resonance experiences an upfield shift indicating a loss of interaction as the phosphate starts to become protonated (Fig. 6b). Under the standard conditions (*T* = 5 °C, pH 6.5–7) the Arg Hη–Nη and Lys Hζ–Nζ were not visible in ^1^H–^15^N HMQC spectra, although at lower pH,^45^ the Lys Hζ–Nζ correlation could be observed and for **1** was slightly downfield in comparison to **3** (Fig. 6c). In addition, ^1^H–^13^C HSQC and ^1^H–^1^H TOCSY spectra revealed that the diastereotopic Arg Hδ protons were resolved in a pH dependent manner for **1** alone, indicating a more restricted side chain conformation for this peptide (Fig. 6d–e). For comparison, ^1^H–^1^H TOCSY spectra showing Arg Hε–Hδ correlations for peptides **3**, **5**, **6** and **7** are given in the Supplementary Information (Fig. S7). The two Arg Hγ resonances for **1** also become non-equivalent on increasing the pH (Supplementary Information, Fig. S8). However, this effect appears to be related to the helicity of the peptide, rather than interaction with the phosphoryl group, since distinct Hγ resonances are also observed for helical, nonphosphorylated peptides **2** and **9**.

**Figure 6.**
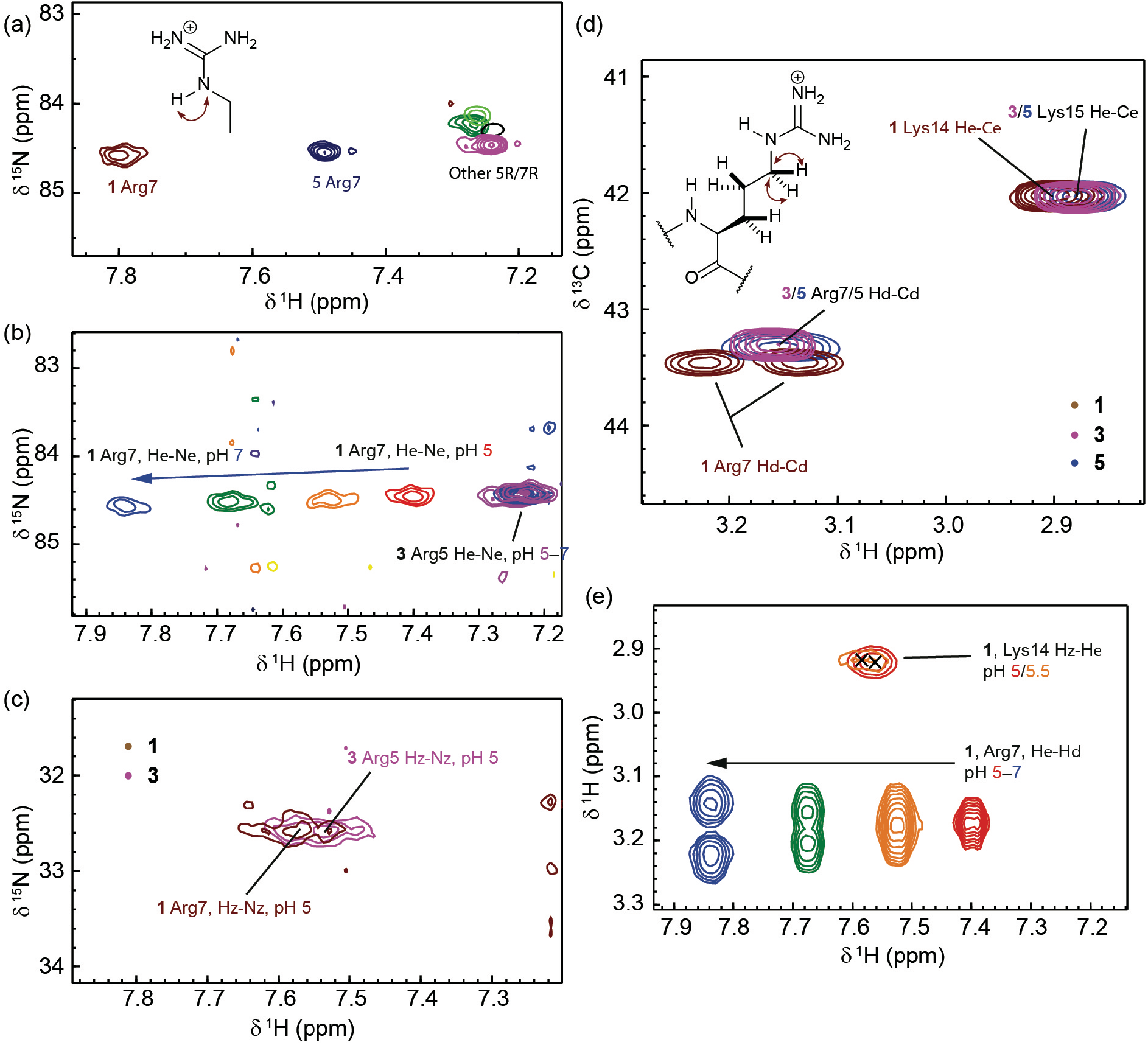
(a) The region of the ^1^H–^15^N HMQC spectra showing Arg Hε–Nε correlations for various peptides at pH 7. (b) The region of the ^1^H–^15^N HMQC spectra showing pH dependent chemical shifts in the Arg Hε–Nε correlation for peptides **1** and **3**. (c) The region of the ^1^H–^15^N HMQC spectra showing a small chemical shift difference in the Lys Hζ–Nζ correlations for peptides **1** and **3** at pH 5.0. (d) The region of ^1^H–^13^C HSQC spectra for peptides **1**, **3** and **5**, pH 7, showing Arg Hδ–Cδ correlations and Lys Hε–Cε correlations. Note the non-equivalence of the two Hδ for peptide **1** only. (e) The region of ^1^H–^1^H TOCSY spectra for peptides **1**, **3** and **5** at pH values 5–7, showing Arg Hε–Hδ and Lys Hζ–Hε correlations. Again, the two Hδ for peptide **1** are not equivalent at high pH. Spectra were recorded on a 600-MHz Bruker Avance spectrometer equipped with a quadruple resonance QCI-P cryoprobe at 5 °C in CD buffer (10 mM NaCl, 1 mM sodium phosphate, 1 mM sodium borate, and 1 mM sodium citrate). The pH was altered by addition of small amounts of 0.1 M HCl or 0.05 M NaOH. Peptide concentrations were in the range 0.5–1.0 mM.

1D ^31^P spectra provided further evidence for differences between peptides in phosphate shielding and altered pSer/pThr side chain dihedral propensities. ^31^P NMR spectra of phospho-peptides contain two peaks: one from the 1 mM free phosphate in the buffer and one from pSer/pThr (Fig. 7). pSer ^31^P resonances appear as a triplet (sometimes unresolved) due to ^3^*J*_PH_ coupling to the two β protons, pThr ^31^P resonances appear as a doublet through coupling to the single β proton. These spectra are highly sensitive to changes in pH. The buffer phosphate peak provides a guide to sample pH through calibration of peak position using standard CD buffer samples at pH 5, 6, 7 and 8. At low pH (pH 5, Fig. 7a), the doublet from peptide **7** appears upfield of the buffer phosphate peak, whereas the triplets from the pSer-containing peptides **1**, **3**, **5** and **6** appear downfield. The ordering of the peptide ^31^P chemical shifts (**7** < **3** < **6** ~ **5** < **1**) at pH 5 follows the helicity. This suggests a de-shielding effect as the phosphate is involved in interactions with more neighbouring basic groups, although the (likely related) effect of altered PO_3_^2−^/PO_3_H^−^ equilibrium could be a more significant factor. At high pH (pH ~ 8, Fig. 7b) the peak positions are all downfield of the buffer phosphate and they are closer together in chemical shift (**7** < **1** < **3** ~ **5** ~ **6**). The fine-structure of the peaks is clearer at pH 8 allowing the ^3^*J*_PH_ coupling constants to be measured (Fig. 7c). There is an inverse correlation between peptide helicity and ^3^*J*_PH_, with the ahelical peptide **7** having the highest ^3^*J*_PH_ (9 Hz) and peptide **1** having the lowest (5.3 Hz). These values match very well with those found by Pandey *et al*.^36^ for pThr- and pSer-containing peptides; the high value for peptide **7** suggests a well-defined P–O–C– H dihedral and ordered pThr side chain, whereas the lower values for peptides **3**, **5** and **1** are consistent with a greater degree of pSer side chain freedom. The reduced value for **1** compared to **3**, **5** and **6** is interesting and suggests altered dihedral propensities for the most helical pSer-containing peptide.

**Figure 7.**
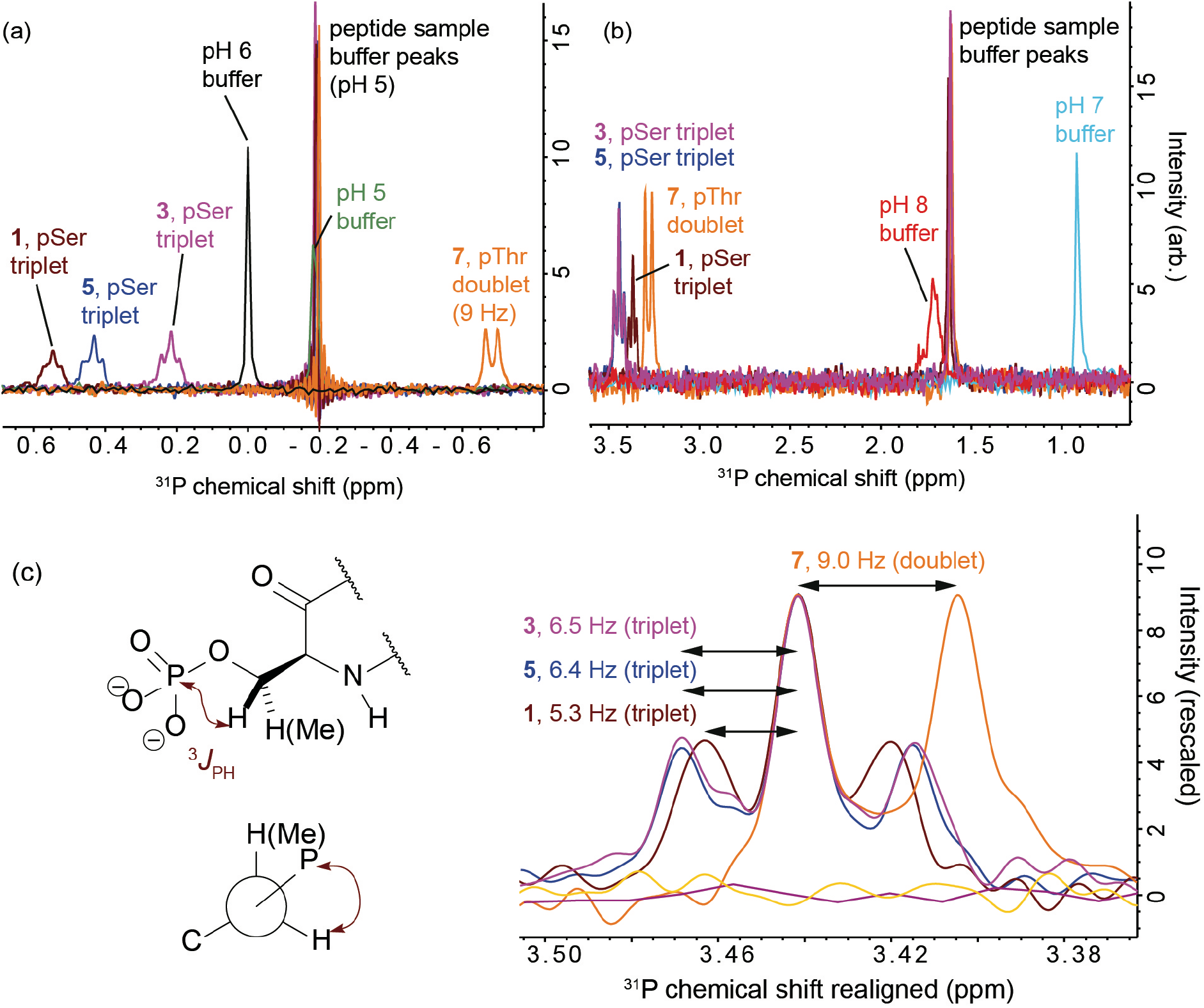
^31^P NMR spectra for selected peptides (**1**, **3**, **5**, **6** and **7**) and CD buffer standards at pH 5, 6, 7 and 8. (a) At pH 5. (b) Just below pH 8. (c) Zoomed-in view of the peak fine-structure at pH ~ 8 with spectra shown realigned to facilitate comparison. Spectra were recorded on a 600-MHz Bruker Avance spectrometer equipped with a quadruple resonance QCI-P cryoprobe at 5 °C in CD buffer (10 mM NaCl, 1 mM sodium phosphate, 1 mM sodium borate, and 1 mM sodium citrate). The pH was altered by addition of small amounts of 0.1 M HCl or 0.05 M NaOH. Peptide concentrations were in the range 0.5–1.0 mM.

## Discussion

We have shown that phosphorylation of Ser residues stabilizes the α-helix in the context of an R_(*i* – 3)_pS_*i*_K_(*i* + 4)_ triad through charge-reinforced side chain interactions with positive co-operativity. This differs to other systems where multiple interactions between side chains are feasible, *e.g*. Glu-Lys-Glu which shows anti-cooperativity,^46^ and provides an exception to the general principle that internal phosphorylation is helix destabilizing. Basic kinase-substrate motifs are common; an Arg at position –3 to the phosphorylated residue is a key selectivity determining residue within the recognition motif of 55 human protein kinases, including Aurora-A and PKA,^47^ and there are over 2000 R_(*i* – 3)_pS_*i*_K_(*i* + 4)_ motifs in human proteins. In contrast, the corresponding Thr variants are not stabilized: Ser is slightly less helix destabilizing than Thr,^4, 5, 11^ whilst the latter favours a β-strand conformation^48^ and branching at β-carbon is helix-destabilizing.^49^ Moreover, recent studies have shown Thr phosphorylation promotes intra-residue hydrogen-bonding between the phosphate and the backbone amide, which would disrupt helix propagation in a manner analogous to the effect of Pro.^36^ Taken together these results highlight the secondary structural sequence context of phosphorylation and highlight the potential for it to affect PPIs not only through orthosteric^13^ or allosteric^50^ changes in recognition features and switching off recognition by changing conformation,^14^ but also through phospho-driven stabilization of an α-helical recognition motif.

## Data Availability

The authors declare that the main data supporting the findings of this study are available within the article and its Supplementary materials. Raw data were generated at the University of Leeds and are available on request from the corresponding author (R.B.).

## Author Contributions

A.J.W. and R.B. conceived and designed the research program, M.B. and R.S.D. designed studies and performed research; M.B. performed NMR and CD analyses, R.S.D. prepared peptides. The manuscript was written by M.B. and R.S.D. and edited into its final form by A.J.W. and R.B. with contributions from all authors.

## Acknowledgements

This work was supported by the EPSRC (EP/N013573/1) and BBSRC (BB/V003577/1). R.S.D. is supported by a studentship from the MRC Discovery Medicine North (DiMeN Doctoral Training Partnership (MR/N013840/1). A.J.W. holds a Royal Society Leverhulme Trust Senior Fellowship (SRF/R1/ 191087). Facilities for NMR spectroscopy and CD spectroscopy were funded by University of Leeds (ABSL award) and Wellcome Trust (108466/Z/15/Z, 094232/Z/10/Z). We thank Nasir Khan and Arnout Kalverda for their support and assistance in this work.

