## Supplementary Information for "α-Helix stabilization by co-operative side chain charge-reinforced interactions to phosphoserine in a basic kinase-substrate motif"

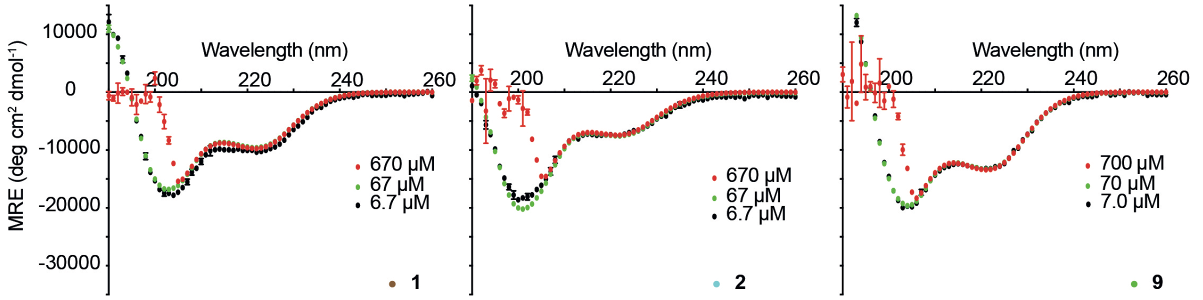


**Figure S1.** The effect of concentration on CD spectra for selected peptides (**1, 2** and **9**). Consistent mean residue ellipticity (MRE) profiles over a range of concentrations indicate peptides are monomeric. At high peptide concentrations, high absorbance leads to inaccuracies in ellipticity measurements below ~207 nm. Experiments were carried out at 5 °C in CD buffer (10 mM NaCl, 1 mM sodium phosphate, 1 mM sodium borate, and 1 mM sodium citrate, pH 7).

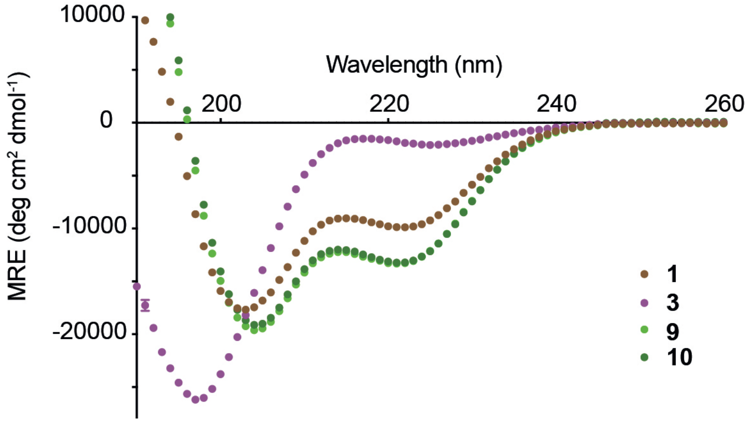


**Figure S2.** CD spectroscopy for model peptides **1**, **3**, **9** and **10**. Experiments were carried out at 5 °C in CD buffer (10 mM NaCl, 1 mM sodium phosphate, 1 mM sodium borate, and 1 mM sodium citrate, pH 7). Peptide concentrations were in the range 50–100 μM.

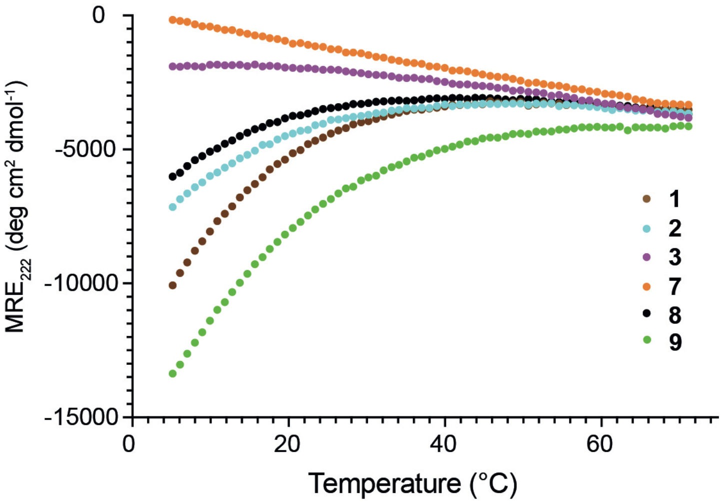


**Figure S3.** Temperature dependent CD spectroscopy analyses for selected peptides **1–3** and **7–9** showing the MRE value at 222 nm highlighting a broad transition from part-helical to coil structure. Experiments were carried out in CD buffer (10 mM NaCl, 1 mM sodium phosphate, 1 mM sodium borate, and 1 mM sodium citrate, pH 7). Peptide concentrations were in the range 50–100 μM.

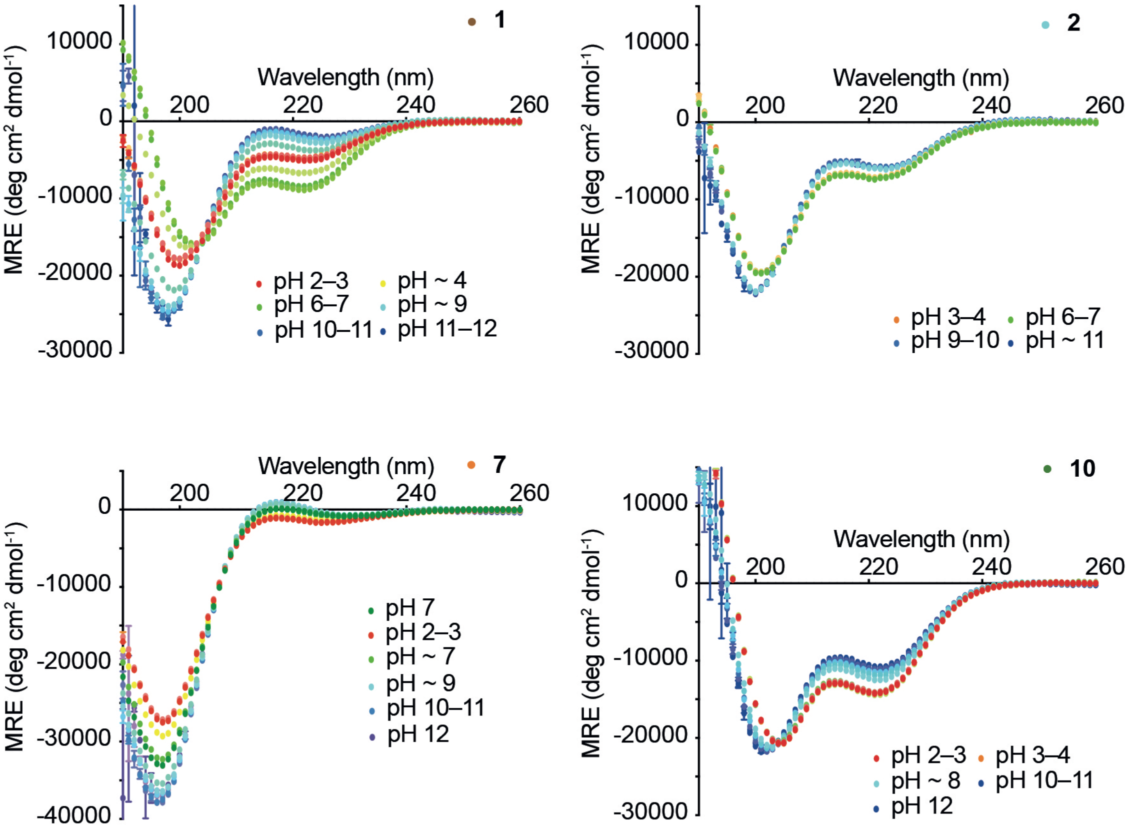


**Figure S4.** The effect of pH on a selection of additional peptides (**2**, **7** and **10**). Spectra for peptide **1** are re-shown from Fig. 3 for comparison. Non-phosphate bearing peptides **2** and **10** exhibit a small loss in helicity at high pH and maintain similar structure under low pH and neutral conditions. The pThr containing peptide **7** is ahelical (<2%) across neutral and alkaline pH range and shows a small amount of helicity (~5%) at low pH, suggesting that singly-charged $PO_3H^-$ is slightly less destabilizing than fully deprotonated $PO_3^{2-}$. Experiments were carried out at 5 °C in CD buffer (10 mM NaCl, 1 mM sodium phosphate, 1 mM sodium borate, and 1 mM sodium citrate). Peptide concentrations were in the range 50–100 μM. The pH was altered by addition of small amounts of 0.1 M HCl or 0.05 M NaOH, with volume changes factored-in to concentration and MRE calculations.

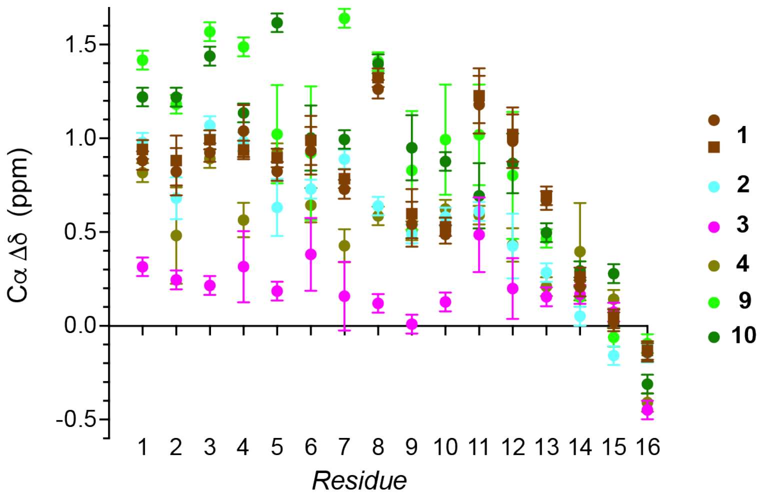


**Figure S5.** Residue specific Cα secondary shifts at 5 °C and pH 6.5–7 are plotted for each residue for peptides **4**, **9** and **10**. For comparison, secondary shifts for peptides **1**, **2** and **3** are repeated from Figure. 5.

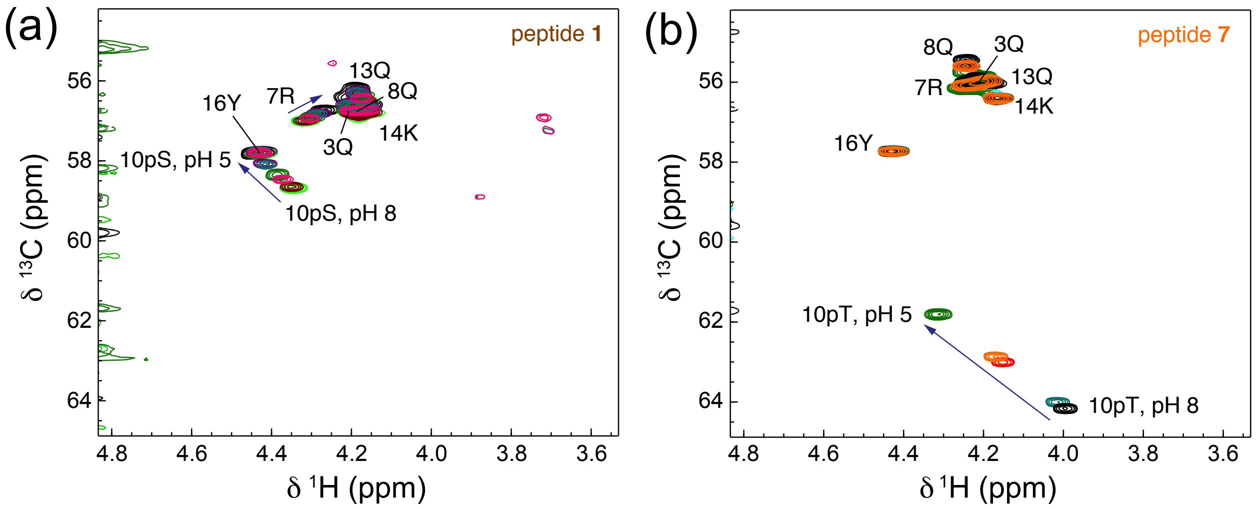


**Figure S6.** Parts of the $^1$H–$^{13}$C HSQC spectra for peptides **1** (a) and **7** (b) at different pH. As predicted from published values,[1] the pSer and pThr Hα–Cα peak changes position with pH. However, as shown in (b), the pThr Cα shift values are higher than expected for a residue in a random coil configuration, indicating that pThr occupies an arrangement unlike that in the QQpTQQ peptide used to generate the values. With the exception of Arg7 for **1**, which shows a subtle shift, other peak positions shown are largely unchanged with pH, and remain appropriate for an unstructured peptide for **7** and a partially helical peptide in **1**. Spectra were recorded on a 600-MHz Bruker Avance spectrometer equipped with a quadruple resonance QCI-P cryoprobe, or a 950-MHz Bruker Ascend Aeon spectrometer equipped with a 5 mm TXO cryoprobe. Experiments were performed in CD buffer (10 mM NaCl, 1 mM sodium phosphate, 1 mM sodium borate, and 1 mM sodium citrate). The pH was altered by addition of small amounts of 0.1 M HCl or 0.05 M NaOH. Peptide concentrations were in the range 0.5–1.0 mM.

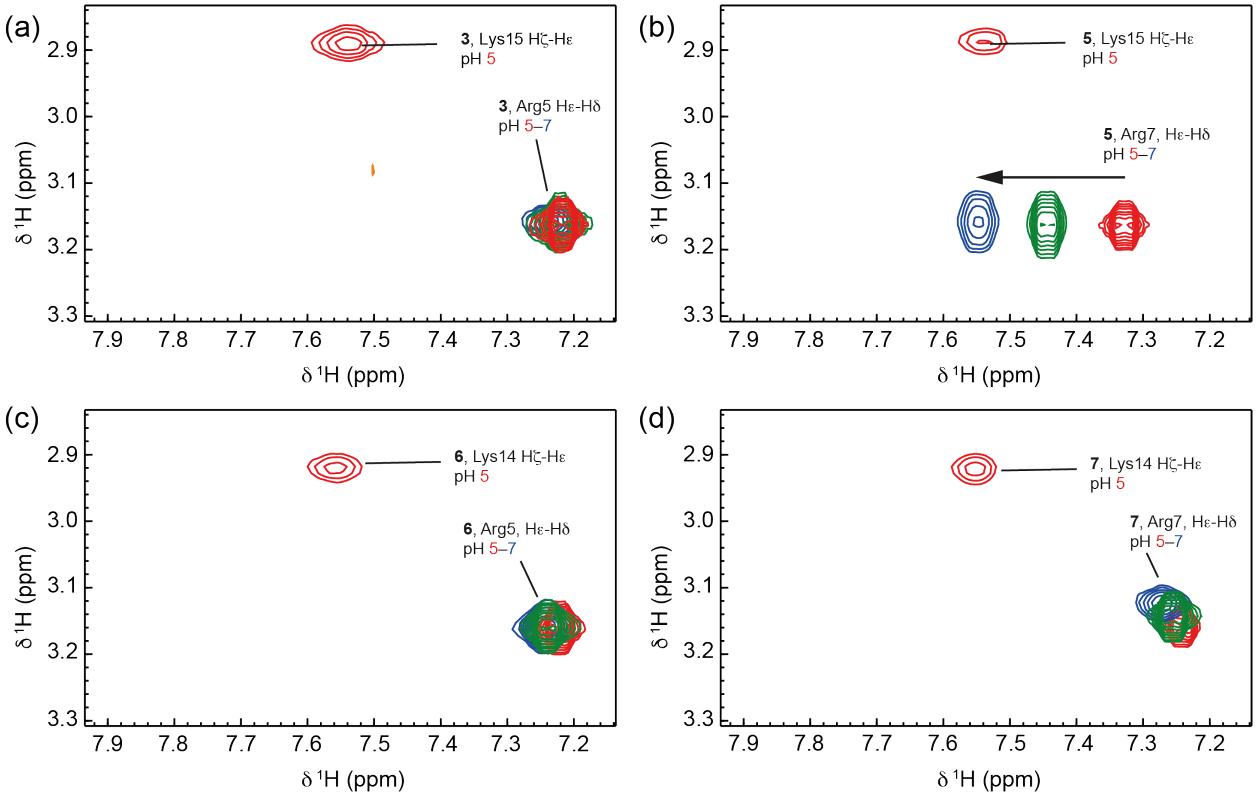


**Figure S7.** The region of $^1$H–$^1$H TOCSY spectra for peptides **3** (a), **5** (b), **6** (c) and **7** (b) at pH 5–7, showing Arg Hε–Hδ and Lys Hζ–Hε correlations. Only peptide **5** shows an effect of changing pH on the Arg Hε shift, and the Arg Hδ have equivalent resonance positions. Spectra were recorded on a 600-MHz Bruker Avance spectrometer equipped with a quadruple resonance QCI-P cryoprobe. Experiments were performed in CD buffer (10 mM NaCl, 1 mM sodium phosphate, 1 mM sodium borate, and 1 mM sodium citrate). The pH was altered by addition of small amounts of 0.1 M HCl or 0.05 M NaOH. Peptide concentrations were in the range 0.5–1.0 mM.

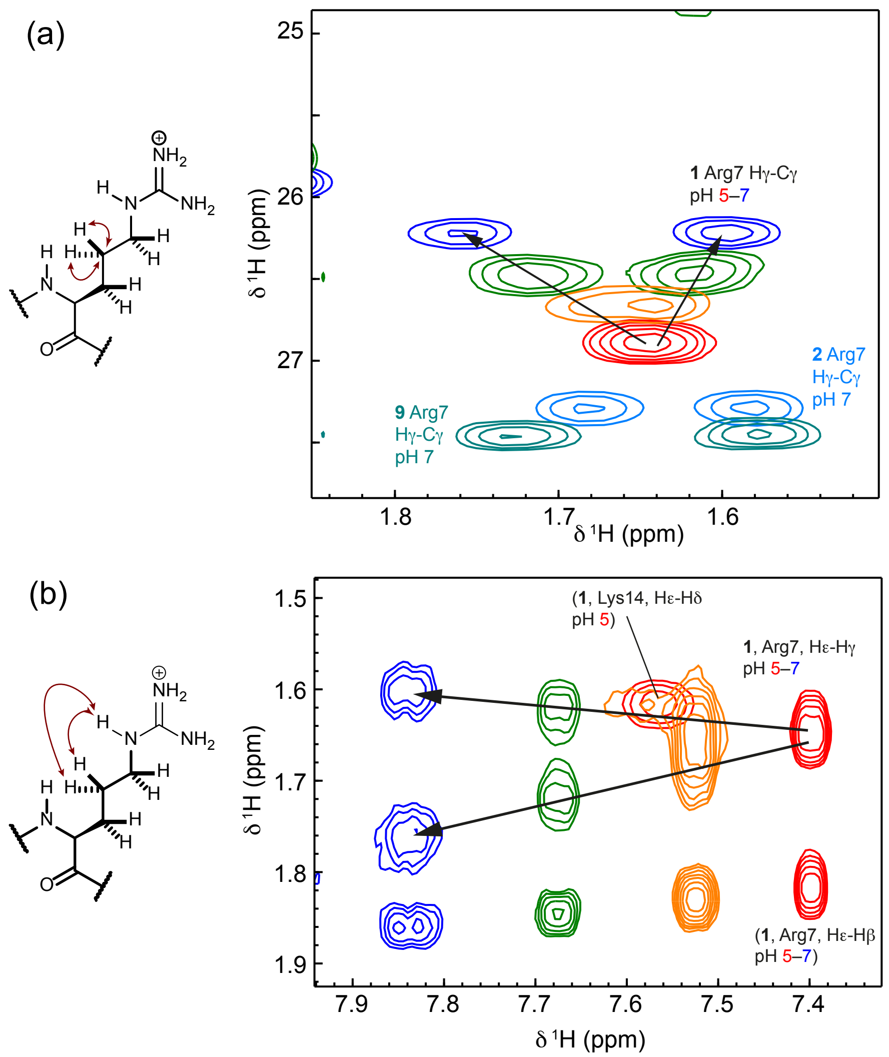


**Figure S8.** (a) The region of $^{1}$H–$^{13}$C HSQC spectra showing Arg Hγ–Cγ correlations for **1** (pH 5–7), **2** and **9** (pH 7). Note the non-equivalence of the two Hγ for peptide **1** at neutral pH as well as for non-phosphorylated but helical peptides **2** and **9**. Low helicity peptides do not show this splitting. (b) The region of $^{1}$H–$^{1}$H TOCSY spectra for peptide **1** at pH values 5–7, showing Arg Hε–Hγ (and Arg Hε–Hβ) correlations. The Lys Hζ–Hδ correlation is also shown at low pH. Again, the two Hγ for peptide **1** are not equivalent at high pH. Spectra were recorded on a 600-MHz Bruker Avance spectrometer equipped with a quadruple resonance QCI-P cryoprobe. Experiments were performed in CD buffer (10 mM NaCl, 1 mM sodium phosphate, 1 mM sodium borate, and 1 mM sodium citrate). The pH was altered by addition of small amounts of 0.1 M HCl or 0.05 M NaOH. Peptide concentrations were in the range 0.5–1.0 mM.

### Peptide Characterization Data

**Table S1**. High resolution mass spectrometry data for peptides

| Peptide | $[M+H]^{1+}$ Obs[d] | $[M+H]^{1+}$ Exp[d] | $[M+2H]^{2+}$ Obs[d] | $[M+2H]^{2+}$ Exp[d] | $[M+3H]^{3+}$ Obs[d] | $[M+3H]^{3+}$ Exp[d] |
|---|---|---|---|---|---|---|
| **1** | 1696.8118 | 1696.8046 | 849.4109 | 849.4023 | 566.6102 | 566.6015 |
| **2** | 1616.8452 | 1616.8383 | 809.4296 | 809.4192 | 539.9541 | 539.9461 |
| **3** | 1696.7538 | 1696.8046 | 849.3837 | 849.4023 | 566.5905 | 566.6015 |
| **4** | 1616.8582 | 1616.8383 | 809.4291 | 809.4192 | 539.9527 | 539.9461 |
| **5** | 1696.7538 | 1696.8046 | 849.3841 | 849.4023 | 566.5905 | 566.6015 |
| **6** | 1696.8105 | 1696.8046 | 849.4108 | 849.4023 | 566.6095 | 566.6015 |
| **7** | 1710.8429 | 1710.8203 | 856.4205 | 856.4102 | 571.2816 | 571.2734 |
| **8** | 1630.9757 | 1630.8540 | 816.4368 | 816.4270 | 544.6259 | 544.6180 |
| **9** | 1600.8658 | 1600.8434 | 801.4322 | 801.4217 | 534.6224 | 534.6145 |
| **10** | 1600.8508 | 1600.8434 | 801.4319 | 801.4217 | 534.6231 | 534.6145 |

Peptide **1**

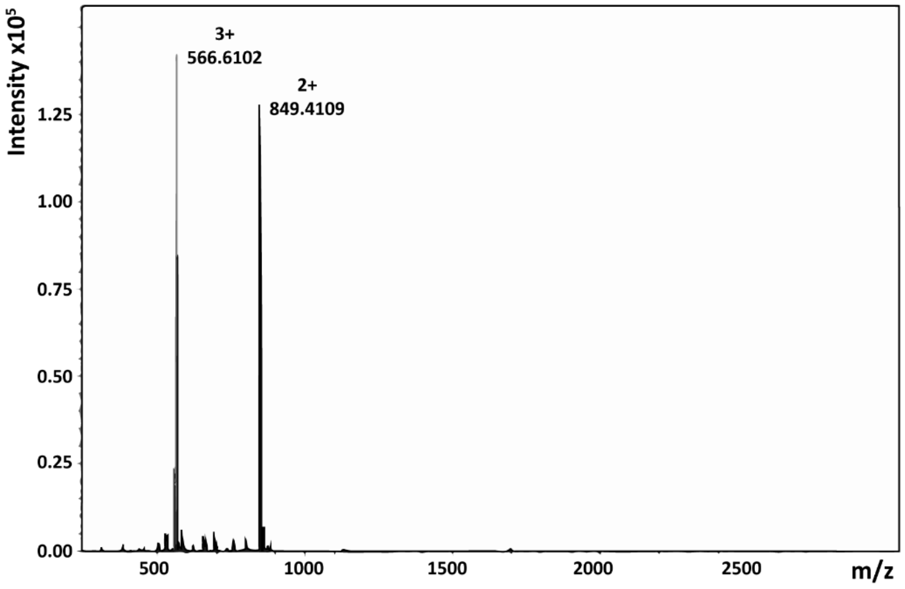

Intensity x10^5
3+
566.6102
2+
849.4109
1.25
1.00
0.75
0.50
0.25
0.00
500
1000
1500
2000
2500
m/z


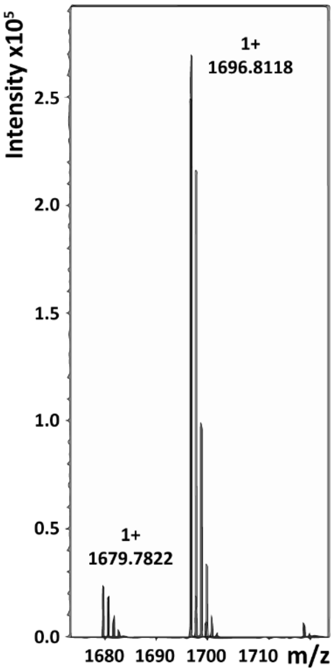

Intensity x10^5
1+
1696.8118
2.5
2.0
1.5
1.0
0.5
1+
1679.7822
0.0
1680
1690
1700
1710
m/z


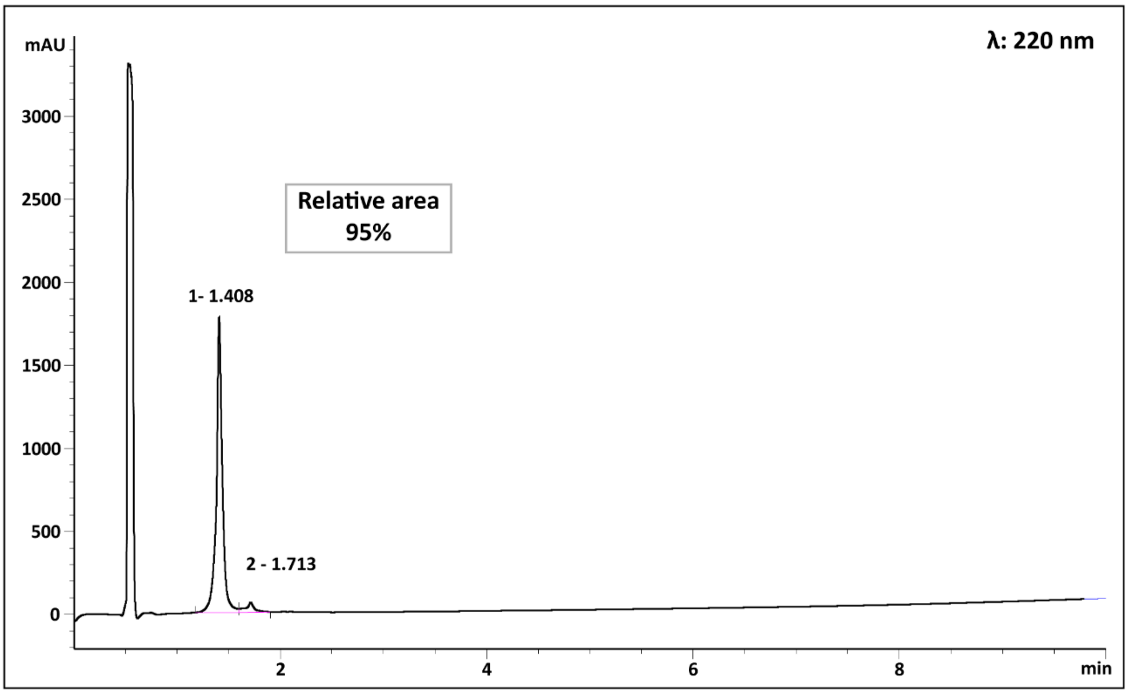

mAU
λ: 220 nm
3000
2500
Relative area
95%
2000
1- 1.408
1500
1000
500
2 - 1.713
0
2
4
6
8
min

Peptide **2**

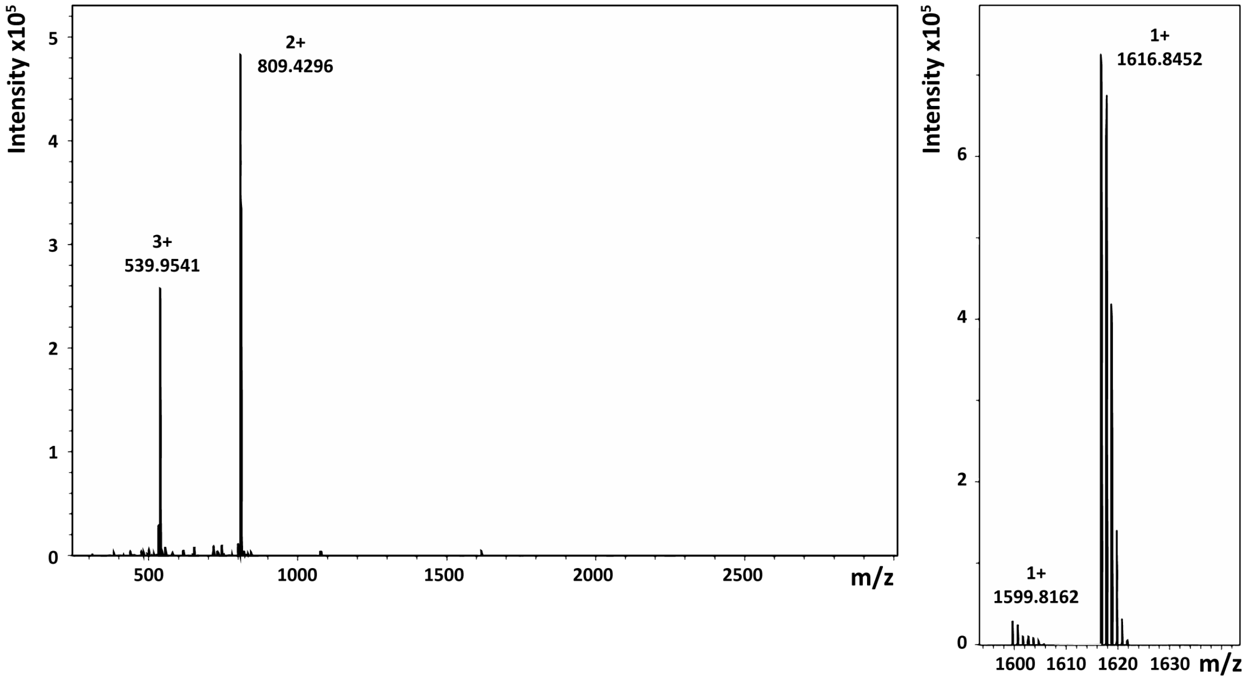
Intensity x10^5
3+
539.9541
2+
809.4296
m/z
1+
1616.8452
1+
1599.8162

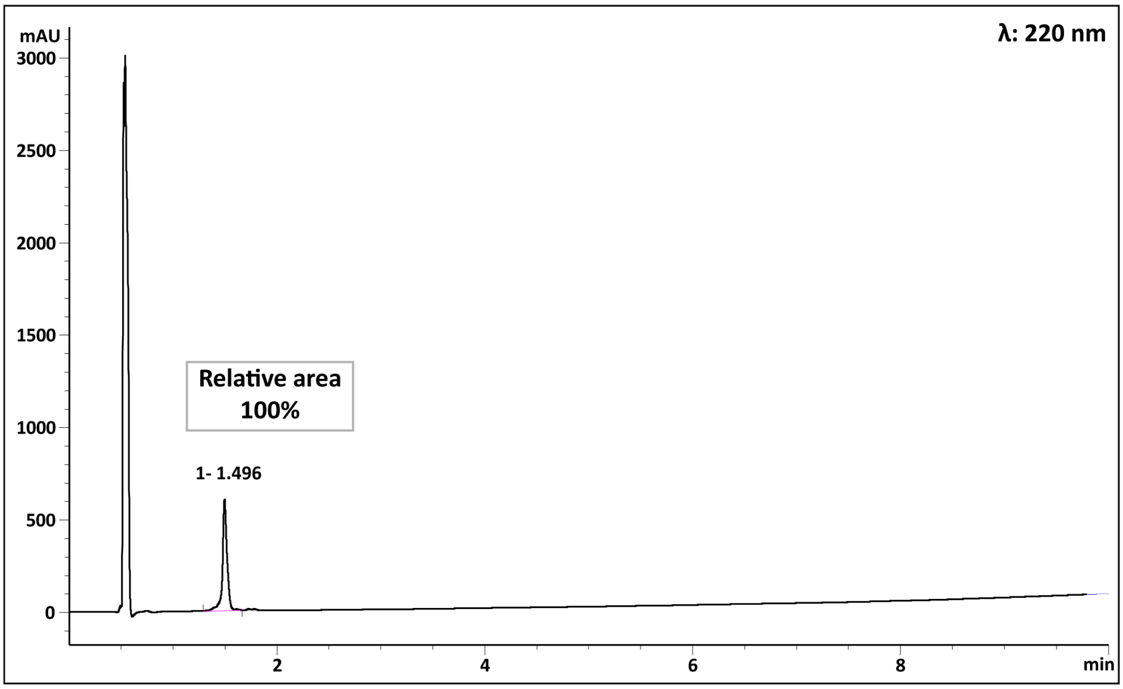
mAU
λ: 220 nm
Relative area
100%
1- 1.496
min

Peptide **3**

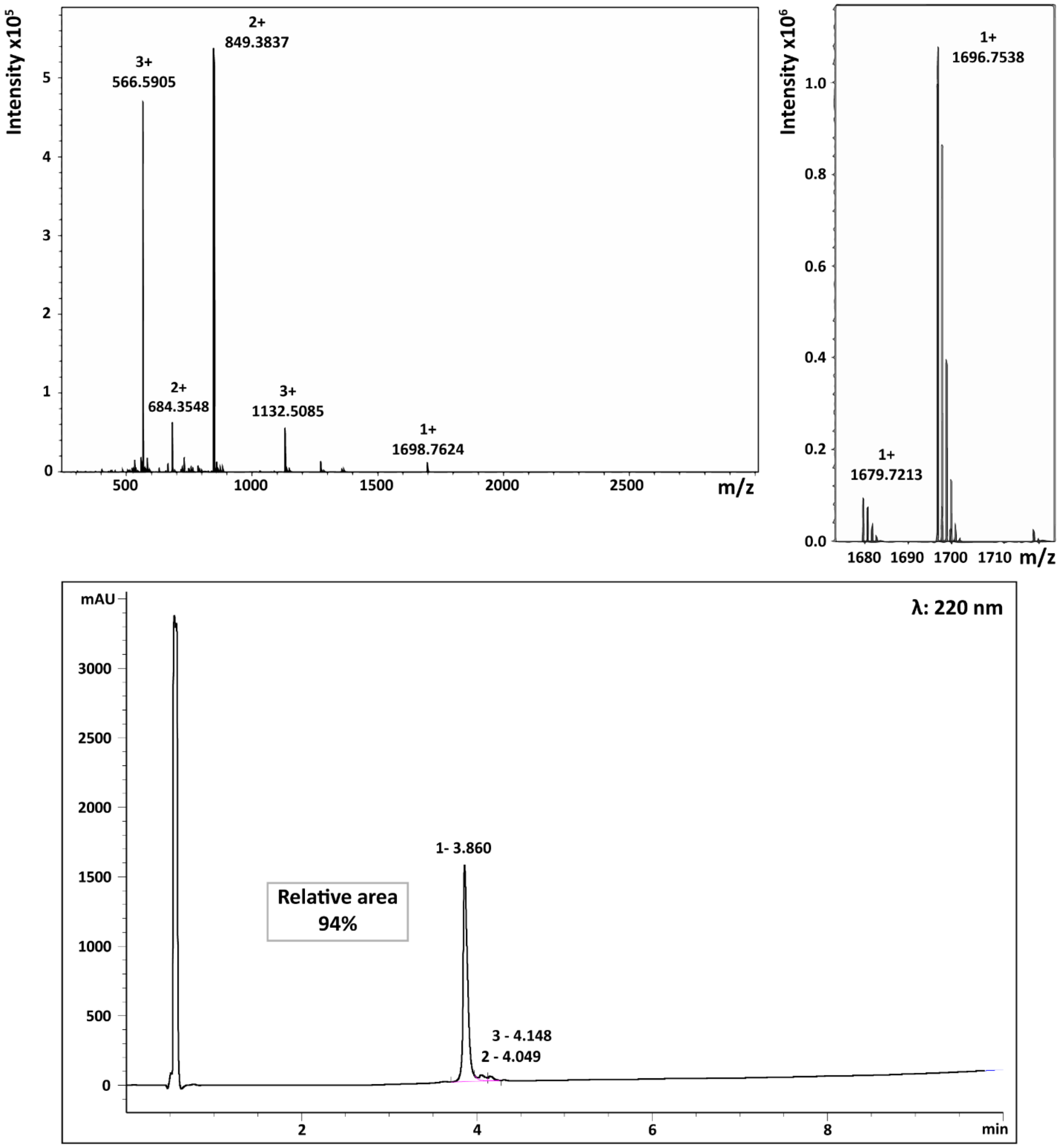
Intensity x10^5
3+
566.5905
2+
849.3837
2+
684.3548
3+
1132.5085
1+
1698.7624
m/z
Intensity x10^6
1+
1696.7538
1+
1679.7213
m/z
mAU
λ: 220 nm
1- 3.860
Relative area
94%
3 - 4.148
2 - 4.049
min

Peptide **4**

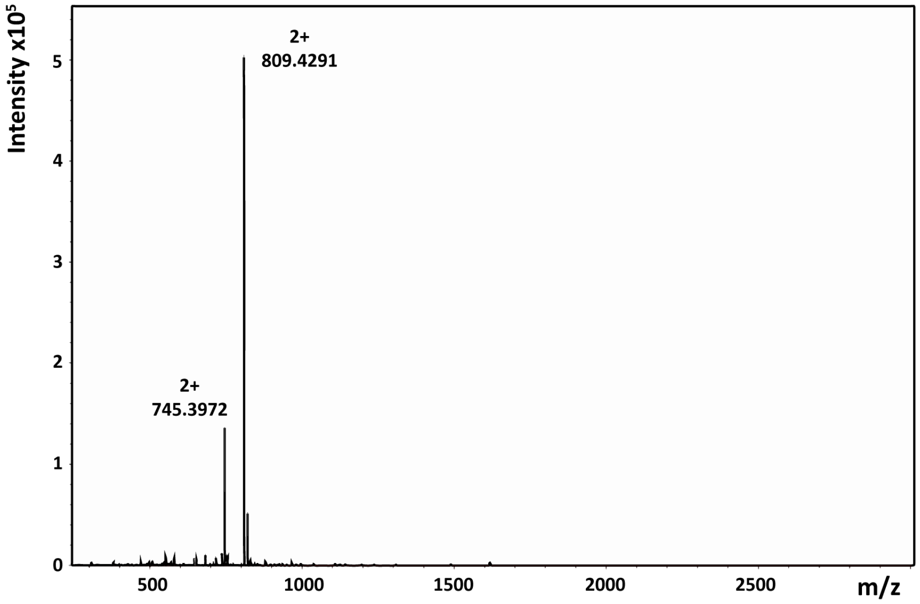

Intensity x10^5
2+
809.4291
2+
745.3972
m/z


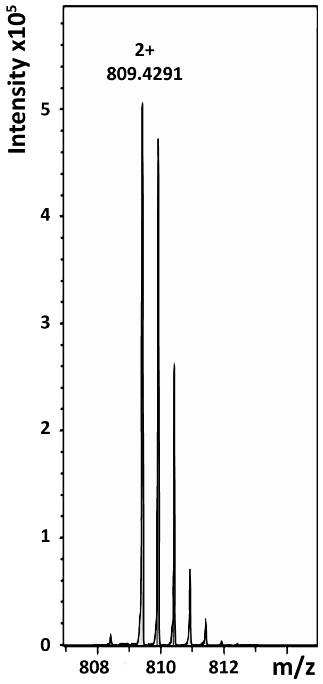

Intensity x10^5
2+
809.4291
m/z


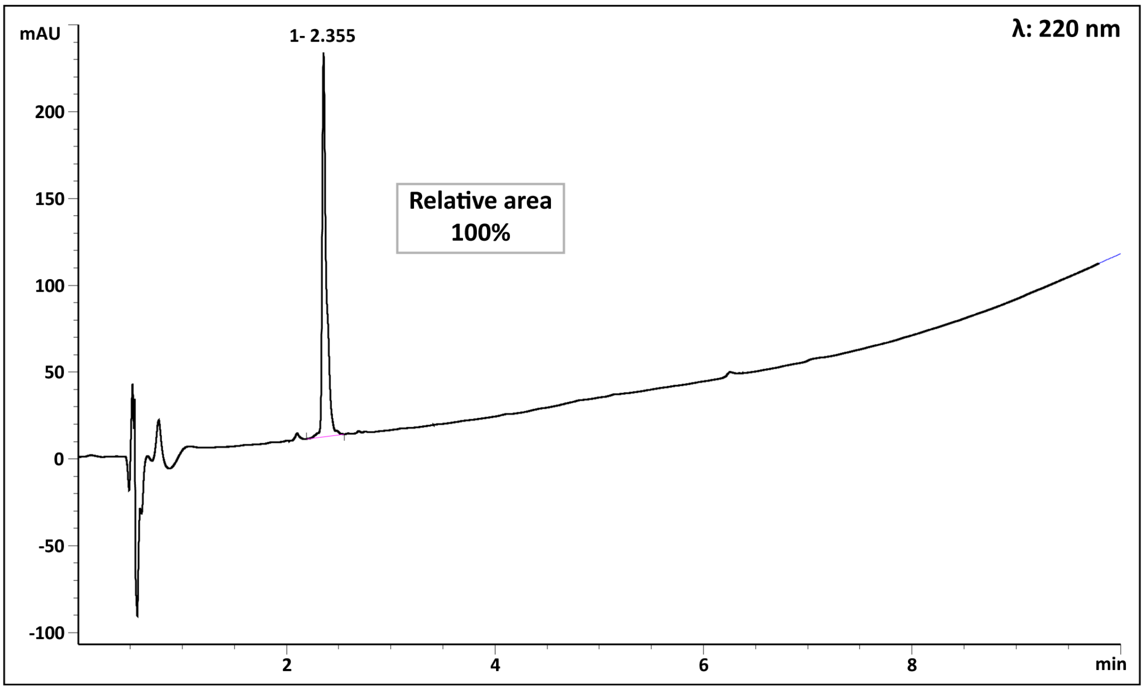

mAU
1- 2.355
λ: 220 nm
Relative area
100%
min

Peptide **5**

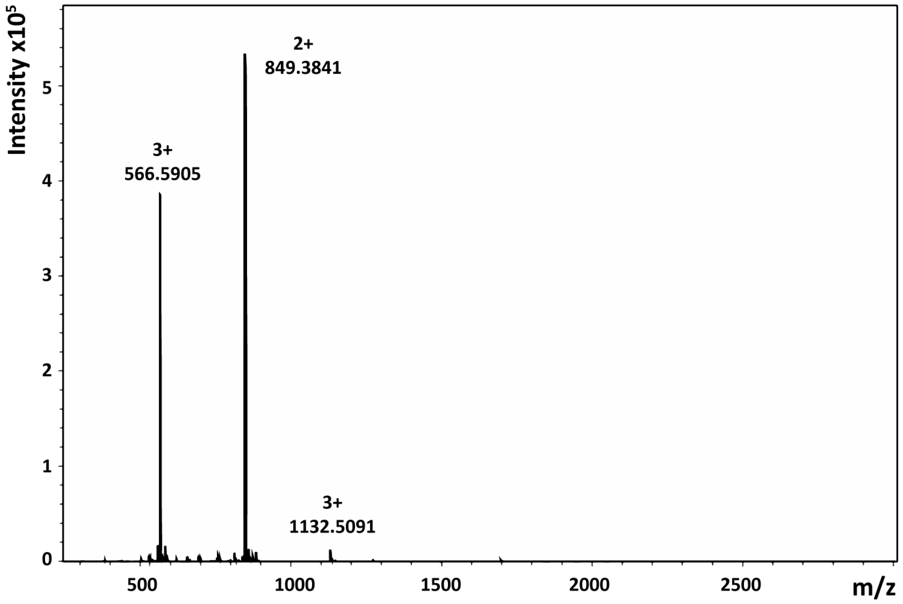

Intensity x10^5
3+
566.5905
2+
849.3841
3+
1132.5091
m/z


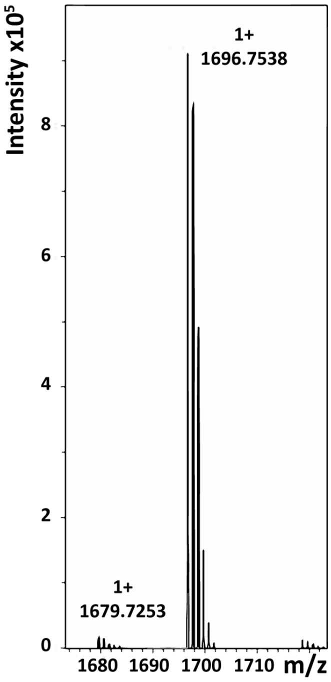

Intensity x10^5
1+
1696.7538
1+
1679.7253
m/z


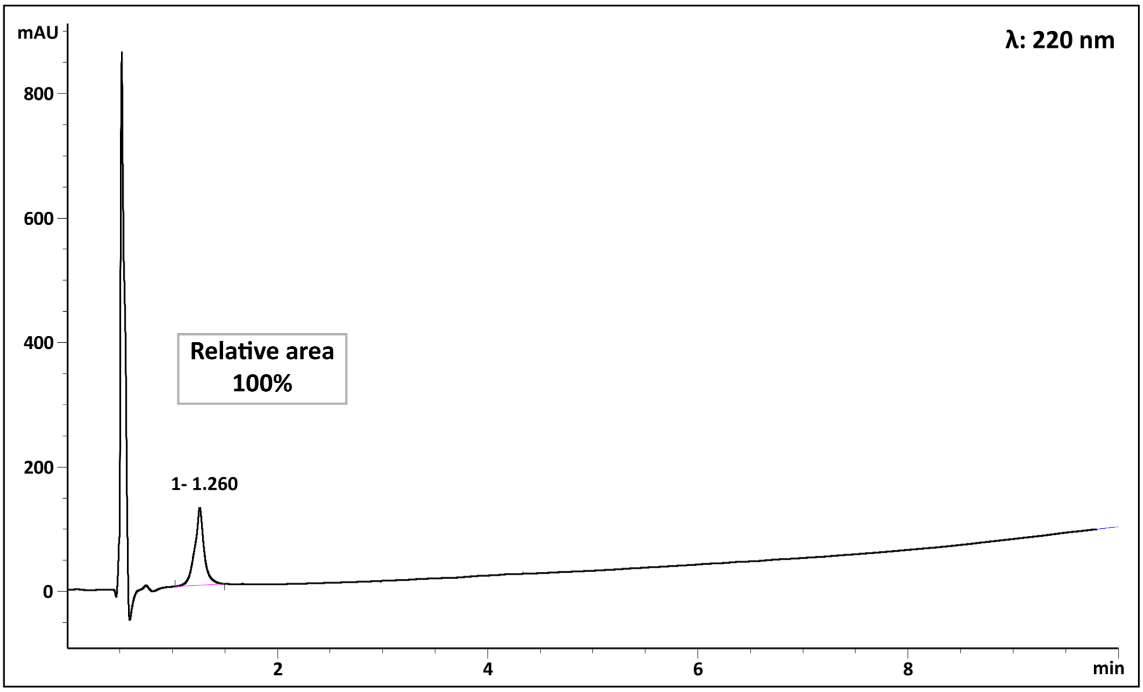

mAU
λ: 220 nm
Relative area
100%
1- 1.260
min

Peptide **6**

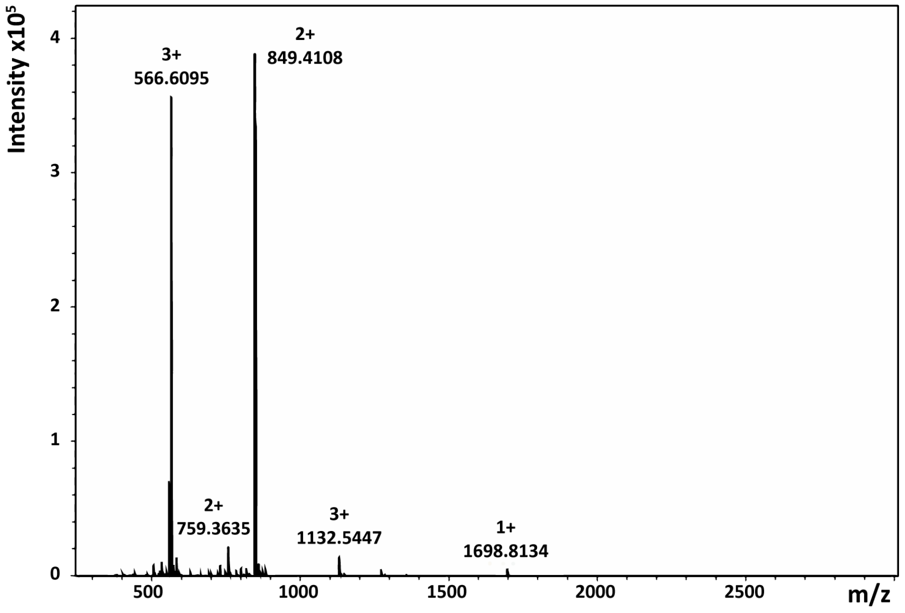
Intensity x10^5
3+
566.6095
2+
849.4108
2+
759.3635
3+
1132.5447
1+
1698.8134
m/z

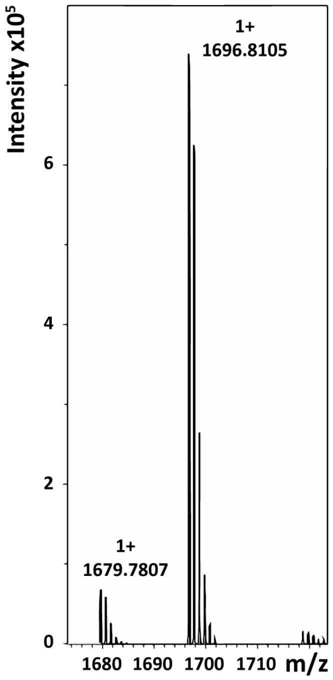
Intensity x10^5
1+
1696.8105
1+
1679.7807
m/z

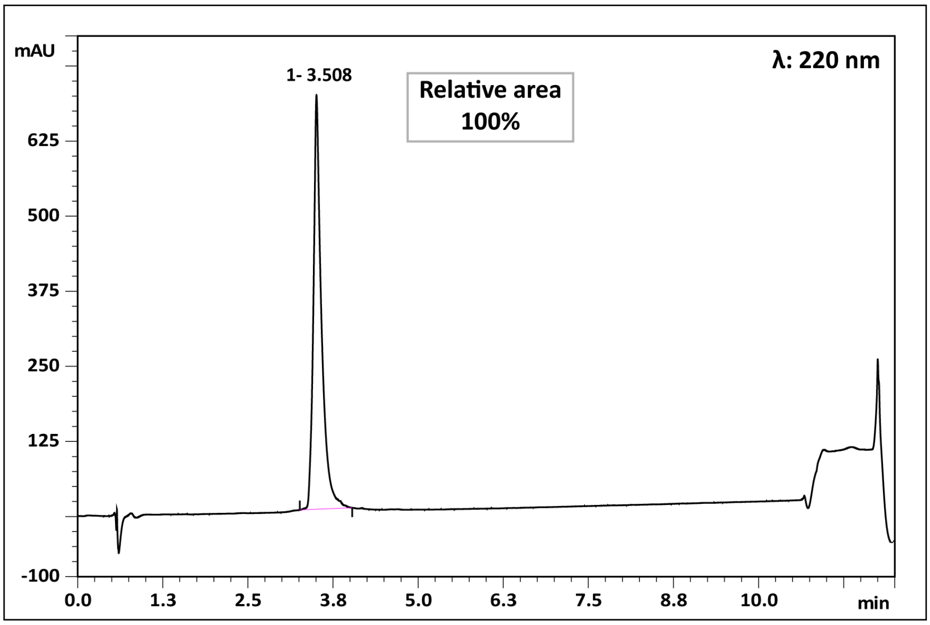
mAU
1- 3.508
Relative area
100%
λ: 220 nm
min

Peptide **7**

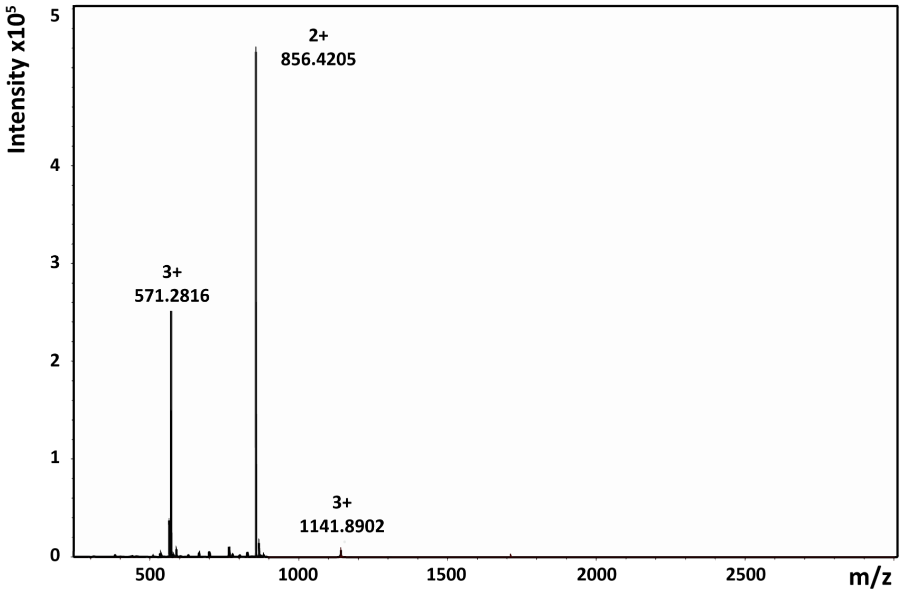
Intensity x10$^5$
2+
856.4205
3+
571.2816
3+
1141.8902
m/z

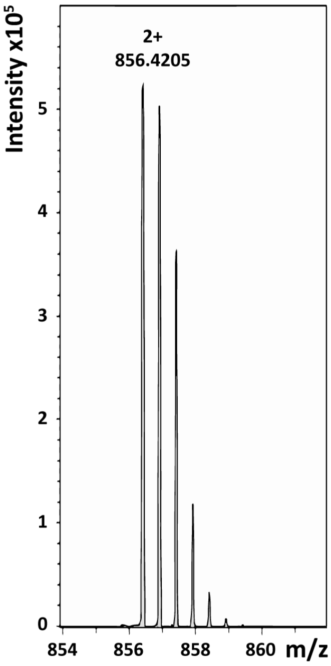
Intensity x10$^5$
2+
856.4205
m/z

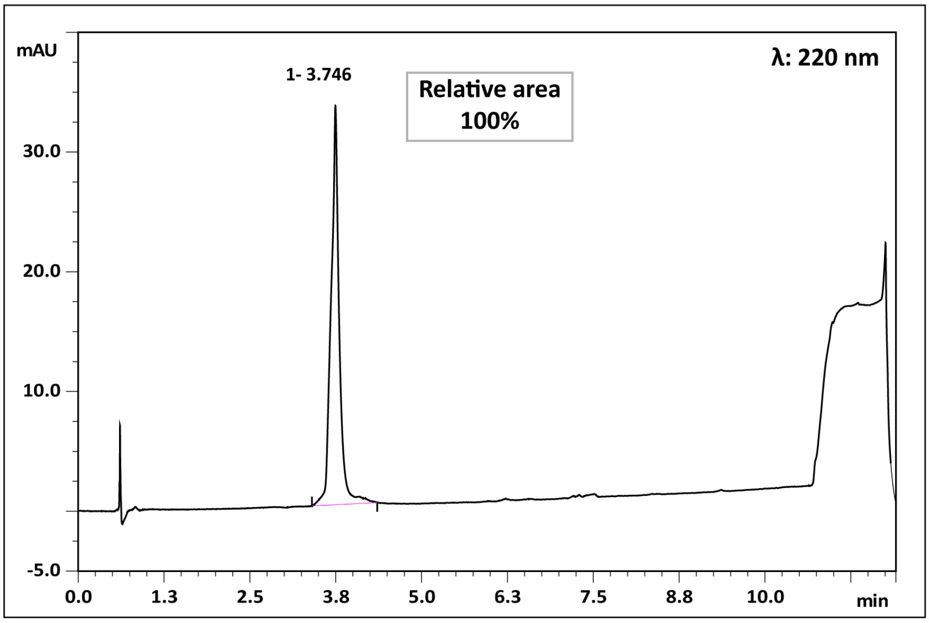
mAU
1- 3.746
Relative area
100%
λ: 220 nm
min

Peptide **8**

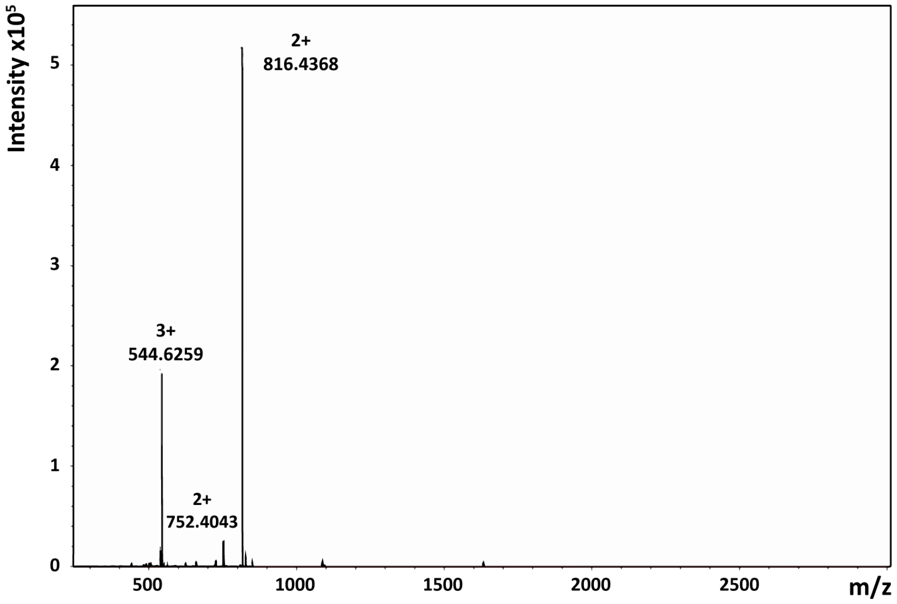
Intensity x10^5
2+
816.4368
3+
544.6259
2+
752.4043
5
4
3
2
1
0
500
1000
1500
2000
2500
m/z

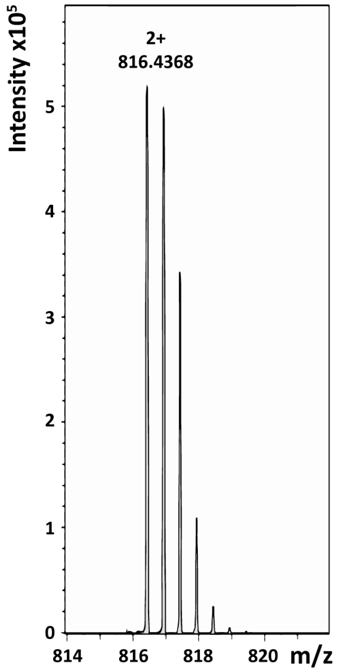
Intensity x10^5
2+
816.4368
5
4
3
2
1
0
814
816
818
820
m/z

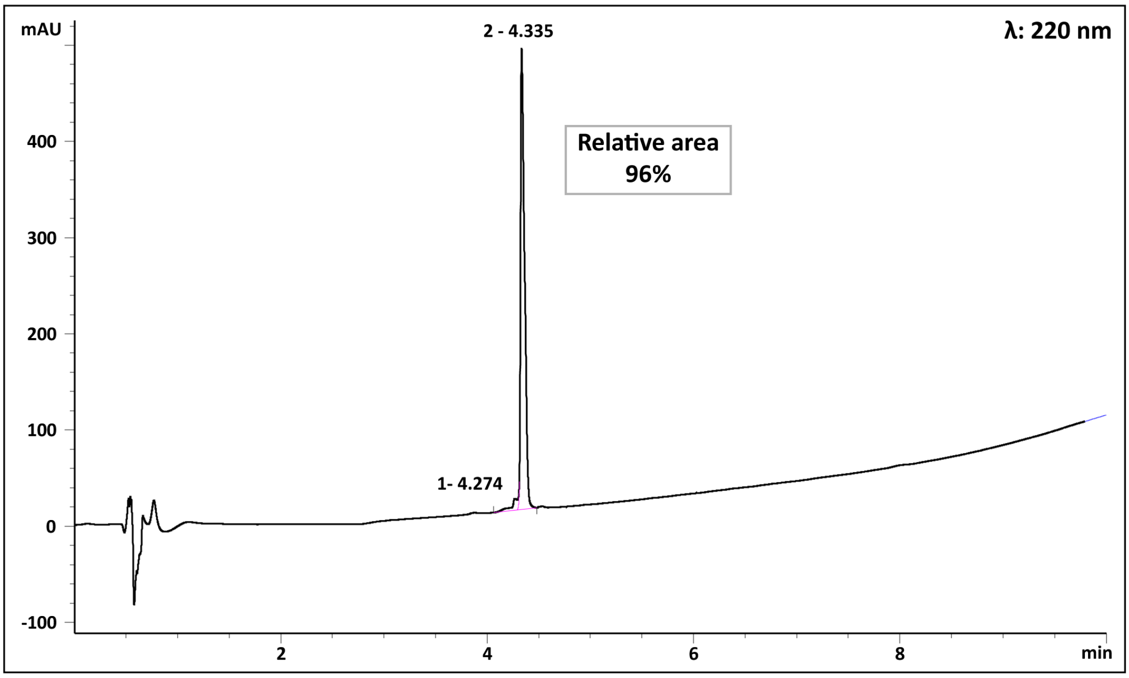
mAU
2 - 4.335
λ: 220 nm
Relative area
96%
400
300
200
100
0
-100
1- 4.274
2
4
6
8
min

Peptide **9**

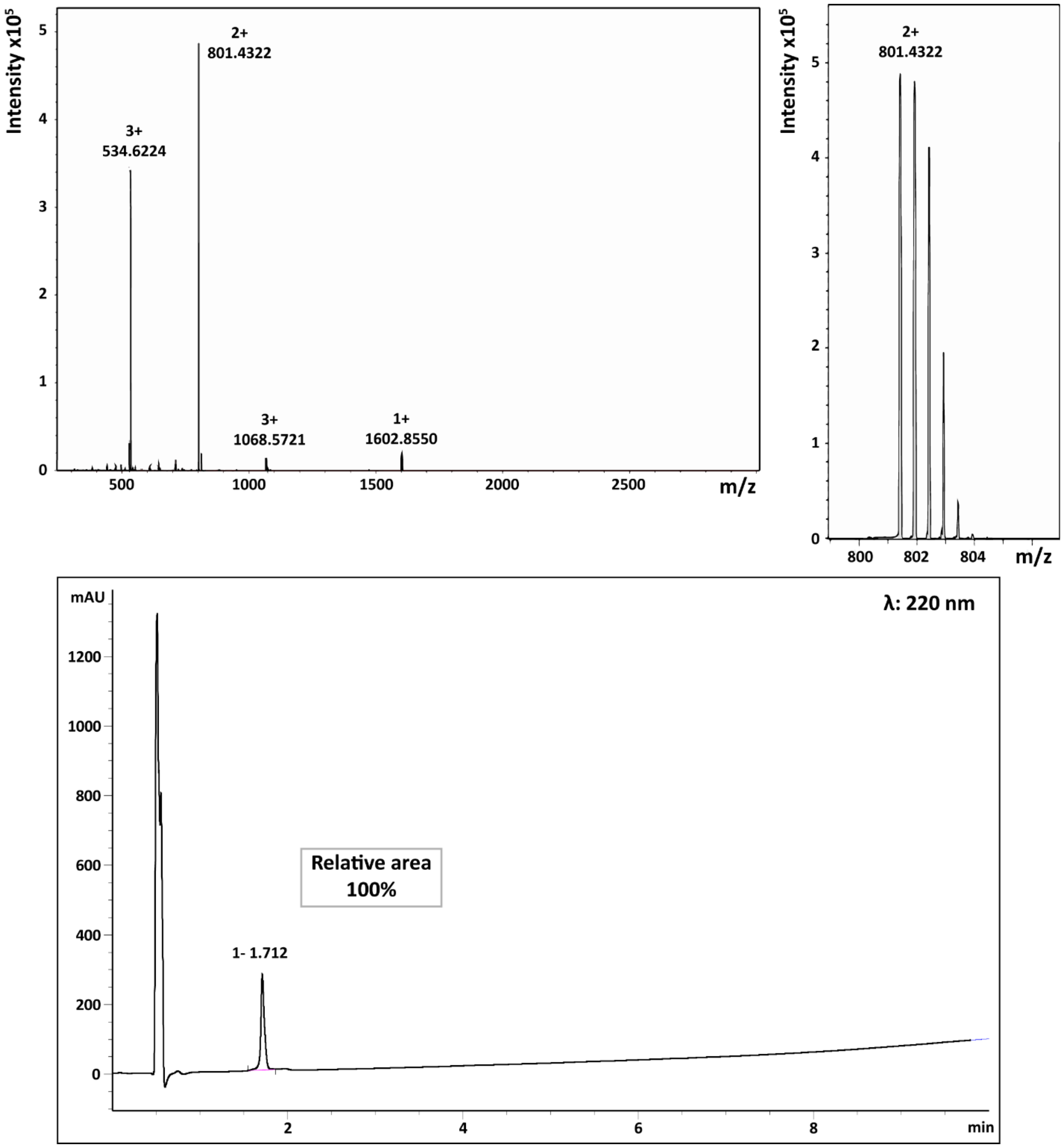
Intensity x10^5
2+
801.4322
3+
534.6224
3+
1068.5721
1+
1602.8550
0
1
2
3
4
5
500
1000
1500
2000
2500
m/z
Intensity x10^5
2+
801.4322
800
802
804
m/z
mAU
λ: 220 nm
1200
1000
800
600
400
200
0
Relative area
100%
1- 1.712
2
4
6
8
min

Peptide **10**

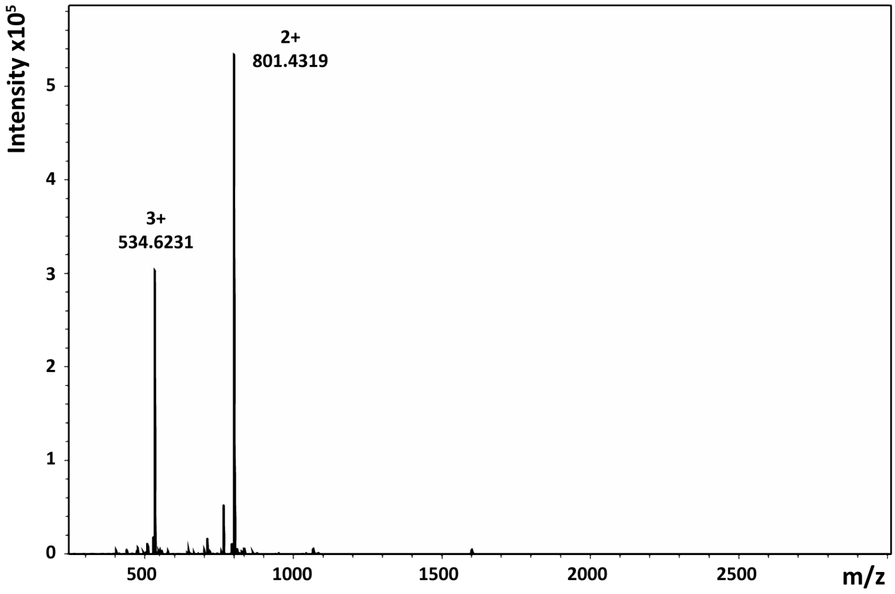


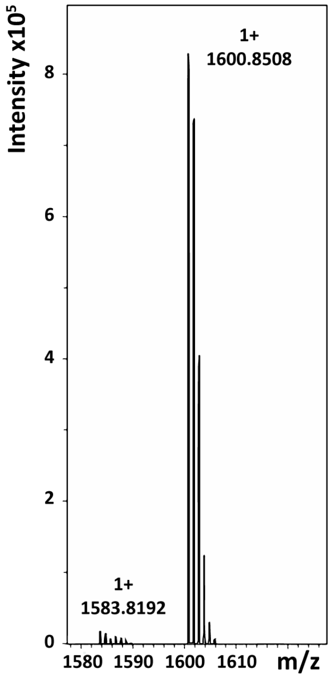


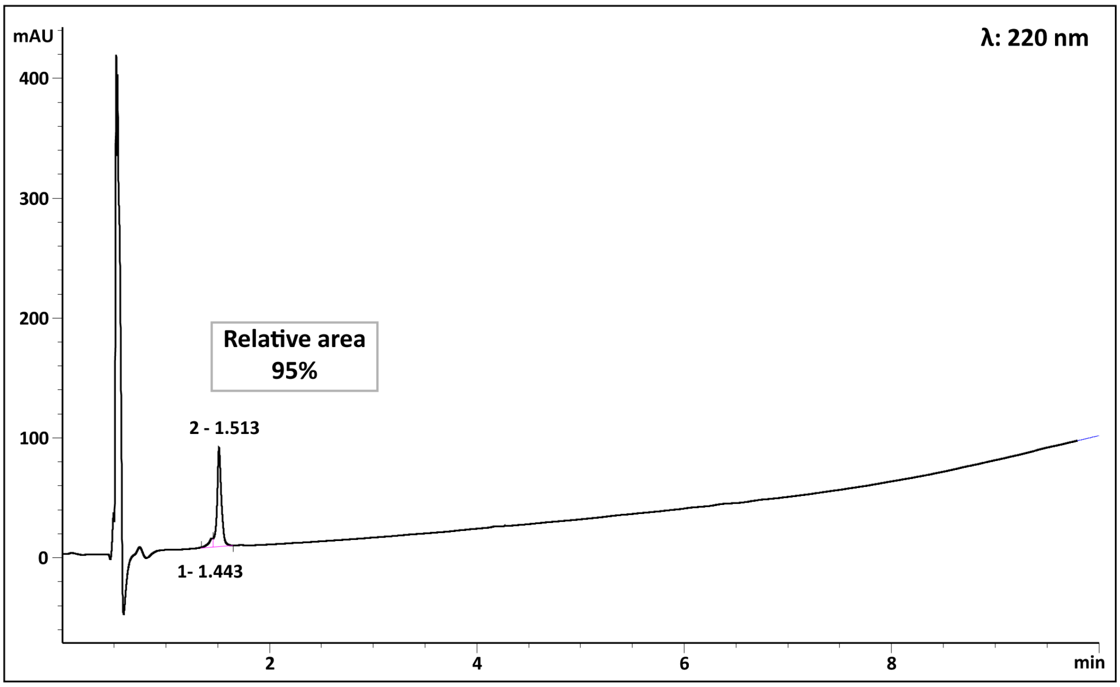
